# Cognitive and neural bases of salience-driven incidental learning

**DOI:** 10.1101/2023.01.23.525123

**Authors:** Sai Sun, Hongbo Yu, Shuo Wang, Rongjun Yu

## Abstract

Humans incidentally adjust their behavioral strategies along feedback dimensions even without explicit reasons to do so. How it occurs may depend on stable individual preferences and contextual factors, such as visual salience. We here examined how task-irrelevant visual salience exerts influence on attention and valuation systems that further drives incidental learning. We first established the baseline behavior with no salience emphasis in Exp.1. We then highlighted either the utility or performance dimension of the chosen outcome using *colors* in Exp.2. We demonstrated that the difference in switching frequency increased along the salient dimension, confirming a strong salience effect. Moreover, the salience effect was abolished when directional information of feedback was removed in Exp.3, suggesting that the observed salience effect is specific to directional feedback. We then generalized our findings using *text* emphasis in Exp.4 and replicated the non-specific salience effects in Exp.5 with simultaneous eye-tracking. The fixation difference between chosen and unchosen values was enhanced along the feedback-specific salient dimension (Exp.4) but kept unchanged when removing directional information (Exp.5). Moreover, behavioral switching correlates with fixation difference, confirming that salience guides attention and further drives incidental learning. Lastly, our neuroimaging study (Exp.6) showed that the striatum subregions encoded salience-based outcome evaluation, while the vmPFC encoded salience-based behavioral adjustments. The connectivity of the vmPFC-ventral striatum accounted for individual differences in utility-driven, whereas the vmPFC-dmPFC for performance-driven behavioral adjustments. Our results provide a neurocognitive account of how task-irrelevant visual salience drives incidental learning by involving attention and the frontal-striatal valuation systems.

## Introduction

Humans adjust their future behaviors based on the current outcome. These adjustments can be based on the utility of the outcomes (e.g., win or loss), performance (e.g., correct or incorrect choice), or both. One of the well-established behavioral strategies deployed during outcome-based adjustments is to stay with the same option as the current one on the next trial after rewarded or correct feedback but to shift to the alternative one after non-rewarded or incorrect feedback (Cavanagh, Hunt, Afraz, & Rolfs, 2010; Chau, Kolling, Hunt, Walton, & Rushworth, 2014; Cohen & Ranganath, 2007; Rudebeck, Saunders, Prescott, Chau, & Murray, 2013). In other words, humans may adjust their behaviors by following an incidental win (or correctness)-stay loss (or error)-shift (WSLS or CSES) rule even without explicit reasons to do so. Such behaviors may reflect our spontaneous thoughts and intrinsic preference and could be exaggerated by contextual factors. However, it is still unclear why we humans learn about these incidental rules. If we do, how do we learn? Under the hypothesis that incidental learning or behavioral adjustments may automatically involve the attention and subjective valuation system, we here examined the behavioral and neural bases of how task-irrelevant visual salience exerts influence on outcome evaluation and further guides irrational decisions.

The salience-driven valuation and decision may reflect how bottom-up visual attention interplays with the internal system by integrating both habitual and goal-directed learning processes. The deployment of attentional gain selectively emphasizes forward connections and links with internal beliefs to plan the next move (Itti & Koch, 2001; Parr & Friston, 2019). Specifically, emphasizing a specific aspect of outcomes increases the behavioral switching along the salient outcome dimension, even if such salient information is redundant (Sun & Wang, 2020; Sun, Yu, & Wang, 2020). The ventral striatum and ventromedial prefrontal cortex (vmPFC) are widely known for their roles in value-based outcome evaluation and action selection, which further guide goal-directed and habitual decisions (Bartra, McGuire, & Kable, 2013; Gläscher, Hampton, & O’Doherty, 2009; Lebreton, Jorge, Michel, Thirion, & Pessiglione, 2009; Lim, O’Doherty, & Rangel, 2011; Rangel & Hare, 2010). The anatomical connectivity within the frontostriatal (vmPFC and medial striatum) circuit accounted for the individual differences in goal-directed and habitual reinforcement learning (Piray, Toni, & Cools, 2016). One recent meta-analysis demonstrated a specific role of the medial prefrontal cortex (mainly the vmPFC) in goal-directed learning, the dorsal striatum in habitual learning, and the ventral striatum in both types of learning (Huang, Yaple, & Yu, 2020). Building on these, we hypothesized that salience might modulate attentional deployment during outcome evaluation and further guide behavioral adjustments by actively recruiting the striatum and vmPFC.

The striatum, marked with dissociable roles in ventral and dorsal portions, has been implicated in salience processing over a range of human neuroimaging and non-human intracranial recording studies (Cooper & Knutson, 2008; Zaehle et al., 2013; Zink, Pagnoni, Chappelow, Martin-Skurski, & Berns, 2006; Zink, Pagnoni, Martin-Skurski, Chappelow, & Berns, 2004; Zink, Pagnoni, Martin, Dhamala, & Berns, 2003). Our previous studies have shown that salience modulates the feedback-related negativity (FRN) and P300 in response to the feedback (Sun & Wang, 2020; Sun et al., 2020). Previous studies also found that the FRN was located in the rostral anterior cingulate cortex (a region close to vmPFC) (Nieuwenhuis et al., 2005; Walsh & Anderson, 2012). The P300 was shown to be related to the ventral striatum BOLD response (Pfabigan et al., 2014). These source localization findings implicate striatum and vmPFC in salience-sensitive feedback processing. Taken together, these findings provide indirect evidence linking the striatum and vmPFC with salience-modulated valuation and decision-making.

To directly examine these links, we employed a simple gambling task with two options and quantified the frequency that participants switched to an alternative option after observing the feedback. Building on our prior EEG studies that employed *text* emphasis for the salience modulation (Sun & Wang, 2020; Sun et al., 2020), in this study, we further examined the attentional deployment and frontal-striatal connectivity during the salience modulation process in separate experiments. Critically, we highlighted either the utility (win or loss) or performance (correct or incorrect) dimension of the *chosen* outcome and we found that the difference in switching frequency was enlarged along the highlighted dimension. When using non-specific salience emphasis of the outcome (i.e., only the utility or performance dimension was emphasized but not the specific outcome), such salience effect was abolished. Moreover, the salience-guided behavioral pattern could be explained by the varied fixation difference between chosen and unchosen values under specific but not non-specific salience manipulation. Using functional magnetic resonance imaging (fMRI), we found that the subregions of striatum were implicated in salience-based outcome evaluation. The vmPFC was implicated in salience-based behavioral adjustments. The vmPFC was functionally connected with the striatum (emphasis on utility) or dmPFC (emphasis on performance), and their connectivity accounted for the interindividual difference in salience-guided outcome-specific behavioral adjustments.

## Materials and Methods

The procedures adopted here closely followed our previous work (Sun & Wang, 2020; Sun et al., 2020), we therefore largely reproduced the methods here, noting variations as appropriate. The major differences between the prior work and the current study are (1) ***replication of Exp.1*** after adding more participants, (2) ***different salience manipulation*** and **behavioral generalization** (color (Exp.2&3&6) vs. *text* (Exp.4&5)), and (3) ***salience-drive attention deployment*** using eyetracking (Exp.4&5) and (4) ***salience-drive neural correlates*** using fMRI (Exp.6). This study was not preregistered.

In Exp.1, we established the baseline behavior by using no salience emphasis which was reported in our prior work (Sun & Wang, 2020; Sun et al., 2020). In Exp.2, we used informative color emphasis that highlights a specific dimension of the chosen outcome, e.g., correct or incorrect. In Exp.3, we used uninformative color emphasis that only highlights the dimension to attend to without directional information, e.g., performance dimension. In Exp.4 and 5, we replicated our behavioral findings in Exp.2 and Exp.3 using informative and non-informative text emphasis, respectively, and further demonstrated the salience-guided attention deployment with simultaneous eye movements recording. In Exp.6, we delineated the neural correlates of salience-guided outcome evaluation and behavioral adjustments using fMRI.

### Participants

There were twenty-one participants (9 male; mean age ±SD: 22.46±1.83 years) who participated in Exp.1 (Behavioral study with *no* emphasis). Thirty participants (5 male; 21.53±2.24 years) participated in Exp.2 (Behavioral study with *specific color* emphasis). Twenty-eight participants (11 male; 20.89±2.45 years) participated in Exp.3 (Behavioral study with *non-specific color* emphasis). Sixty-two participants were recruited for Exp.4 (Eye-tracking study with *specific text* emphasis). Thirty-eight participants were recruited for Exp.5 (Eye-tracking study with *nonspecific text* emphasis). Fourteen eye-tracking participants from Exp.4 and another six participants from Exp.5 were dropped from further analysis due to the high rejection rate of trials (>17%) including trials in which responses were initiated too quickly (< 100 ms) during the choosing period, and those in which the fixation duration for either of the two cards is less than 100ms. The remaining eye-tracking participants were 48 (17 male; 20.63 ± 2.88 years) for Exp.4 and 32 (15 male; 20.90 ±2.11 years) for Exp.5. Twenty-eight participants participated in Exp.6 (fMRI study with *specific color* emphasis), and 3 of them were dropped from further analysis due to strong head motions (>3 mm, in x, y, and z-axis), leaving 25 participants (8 male; 22.15±2.40 years) in total. All participants had a normal or corrected-to-normal vision, but not any neurological or psychiatric disorders. All participants provided written informed consent according to protocols approved by the South China Normal University Institutional Review Board. All participants gave written, informed consent and were informed of their right to discontinue participation at any time.

### Stimuli and procedure

We employed a well-established paradigm to study the behavioral adjustment (Sun & Wang, 2020; Sun et al., 2020). At the beginning of each trial, participants viewed two gambling cards (rough visual angle 15° × 8°) and were required to choose one card by pressing the corresponding button within 1.5 seconds (**Fig. 1A, B, D**; 2 seconds for fMRI experiment). The trial was discarded if participants did not make a response within 1.5 seconds (2 seconds for the fMRI experiment). The chosen card was highlighted by a yellow box immediately after the button press for the rest of the 1.5 seconds (2 seconds for the fMRI experiment). Subsequently, the outcome associated with both the chosen card and unchosen card was shown for 1.5 seconds (4 seconds for the fMRI experiment), followed by an inter-trial-interval (ITI) of 0.5 seconds (jittered randomly with a uniform distribution between 1 to 2 seconds for fMRI experiment). We used E-prime 2.0 (Psychology Software Tools, Inc. Pittsburgh, PA, USA, www.pstnet.com/e-prime) for stimulus presentation and response recording.

**Fig. 1.**
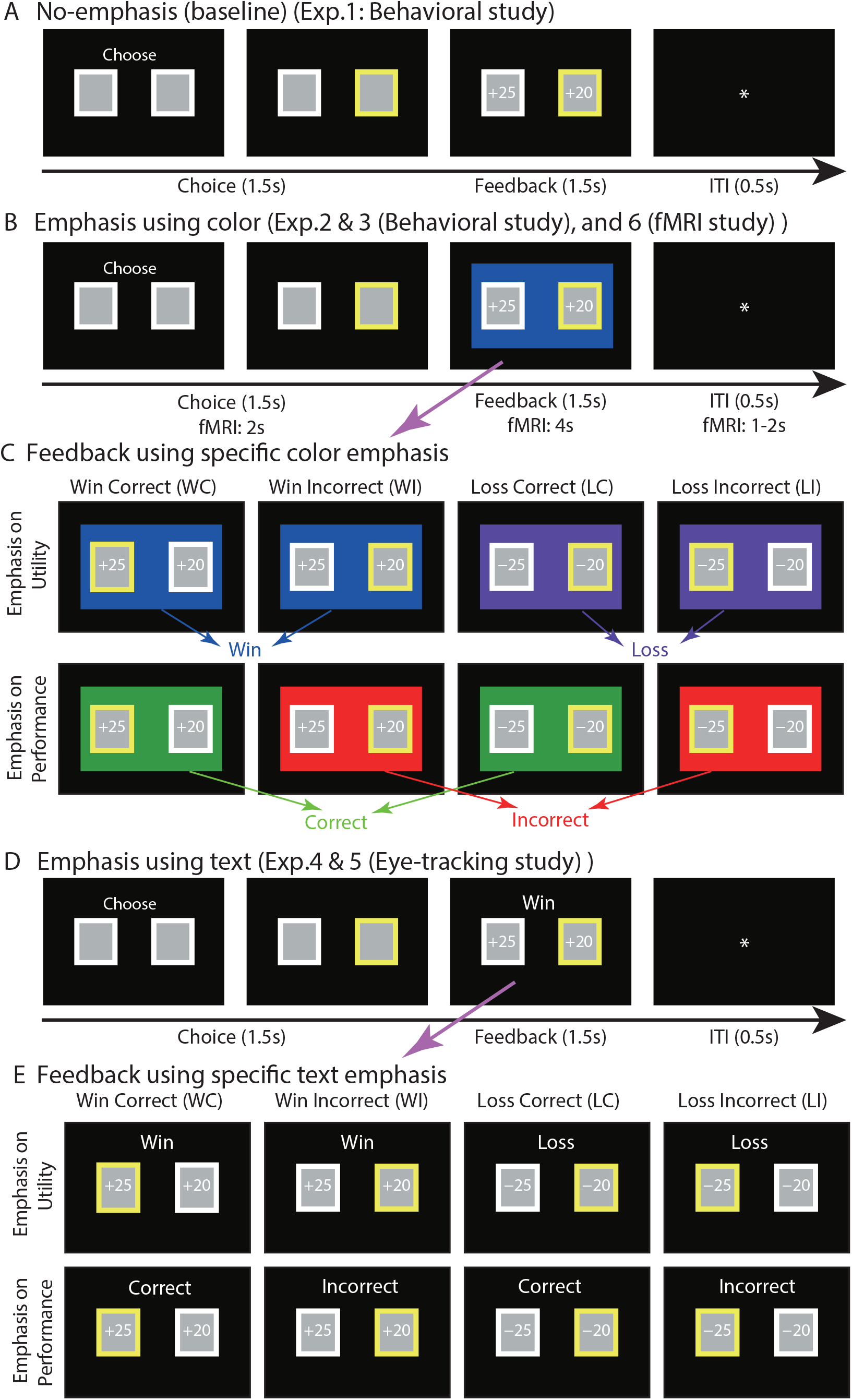
Task and stimuli. **(A)** Task. At the beginning of each trial, two gambling cards were presented and participants were required to choose one card. The chosen card was highlighted in yellow. Then both the outcome associated with the chosen card and the alternative outcome were shown, followed by an inter-trial interval. **(B)** In Exp.2, 3, and 6, a colored rectangle was displayed surrounding the outcomes to emphasize a task aspect. **(C)** Types of outcomes. There were four types of outcomes: win-correct (WC), win-incorrect (WI), loss-correct (LC), and loss-incorrect (LI). “Win” and “Loss” mean that the chosen card yields a (“+”) and penalty (“-”), respectively. “Correct” and “Incorrect” mean that the chosen card yields a better (a larger reward or a smaller penalty) and worse (a smaller reward or a larger penalty) outcome compared to the unchosen card, respectively. In Exp.2, and 6, each color was associated with one specific type of outcome, whereas in Exp.3, the two colors associated with each dimension of emphasis (e.g., red/green for utility and blue/purple for performance) were presented randomly. **(D)** In Exp.4 and 5, a non-colored text was displayed above the outcomes of the cards to indicate the emphasized dimension. **(E)** In Exp.4, participants were explicitly told the association between the texts and chosen outcome (e.g., “+” for Win, “-” for Loss, “a larger reward or a smaller penalty” for Correct, and “a smaller reward or a larger penalty” for Incorrect). In Exp.5, a similar procedure as Exp.4 was performed except that a non-specific outcome message about the emphasized dimension (‘Win or Loss’, ‘Correct or Incorrect’) was displayed above the outcomes.

There were four types of outcomes: win-correct (WC), win-incorrect (WI), loss-correct (LC), and loss-incorrect (LI). ‘Win’ and ‘loss’ mean that the chosen card yields a reward and penalty, respectively (**Fig. 1C, E**). ‘Correct’ and ‘incorrect’ mean that the chosen card yields a better (a larger reward or a smaller penalty) and worse (a smaller reward or a larger penalty) outcome compared to the unchosen card, respectively. Four corresponding examples were given and explicitly explained to each participant (see **Fig. 1C, E** for examples). Unbeknownst to participants, all outcomes were predetermined (the same for all participants) and pseudorandomized across conditions. Each pair of chosen and unchosen cards was presented randomly within each condition. Specifically, the value of the chosen card was randomly decided (in integers) from a uniform distribution ranging from −40¥ to **+**40¥ (about $6.2), whereas the value of the unchosen card was also determined randomly from a uniform distribution, but with the constraint that the absolute difference between the chosen and unchosen outcomes was less than 20¥ (but no less than 2¥; the chosen and unchosen outcomes could have different signs in the same trial). The values of the chosen and unchosen cards were independent of card positions. Participants were told that their goal was to earn as much money as possible, and they were free to employ any strategies to achieve that goal.

Before the experiment, participants were informed that one trial would be selected randomly from the experiment, and the value of the chosen outcome would be added to (or subtracted from) their base payment (60¥ (about $10)). The payoff was given based on whether they won or lost the gamble and varied from −40¥ to **+**40¥ (about $6.2). Ten practice trials were given before the experiment, allowing participants to familiarize themselves with our procedure. No reward was given for practice trials.

### Salience manipulation

Each participant underwent two sessions. In Exp. 1 (**Fig. 1A**), both sessions were the same and had no salience emphasis. This baseline condition has been reported in our previous study (Sun & Wang, 2020; Sun et al., 2020). In Exp.2-6, each session had a different salience manipulation (emphasizing one of the task aspects using either color (**Fig. 1B**, Exp.2, 3, and 6) or *texts* (**Fig. 1D**, Exp.4 and 5). To emphasize utility (win/loss) or performance (correct/incorrect), a colored rectangle (i.e., highlight) was displayed around the outcomes of the cards (**Fig. 1B**), or a noncolored text was displayed above the outcomes of the cards (**Fig. 1D**). In Exp.2, the highlight was specific to the outcome. Participants were explicitly told the association between each color and outcome (e.g., blue for win, purple for loss, green for correct, and red for incorrect; see **Fig. 1C** for examples, corresponding to **Fig. 1B**), and colors were randomly assigned to each outcome across participants. However, in Exp.3, two colors (e.g., red/green for utility and blue/purple for performance) were presented randomly for each session, and participants were told that when red/green was presented, they should pay attention to the utility (i.e., win/loss) dimension of the outcomes, whereas when blue/purple was presented, they should pay attention to the performance (i.e., correct/error) dimension. In contrast to Exp.2, the highlight in Exp.3 only reminded participants which dimension they should focus on without providing any directional information about the chosen outcome.

Except for the color emphasis (perceptual salience), two independent behavioral experiments using *text* emphasis (semantic salience) were performed to quantify the attentional deployment combined with simultaneous eye movements recording in Exp.4 and 5 (**Fig. 1D**). In Exp.4, participants were explicitly told the association between the texts and chosen outcome (e.g., “+” for Win, for Loss, “larger positive value or smaller negative value” for Correct, and “larger negative value or smaller positive value” for Incorrect; see **Fig. 1E** for examples, corresponding to **Fig. 1D**). In Exp.5, a similar procedure as Exp.4 was performed except that a non-specific highlight message about the emphasis dimension (‘Win or Loss’, ‘Correct or Incorrect’) was displayed. Lastly, a similar procedure as Exp.3 (Fig. 1B) was performed to demonstrate the functional role of a frontal-striatal circuit in salience-guided outcome evaluation and behavioral adjustments combined with fMRI.

The colors were partially counterbalanced across participants for Exp.2, 3, and 6 across two sessions (salience emphasis) and outcomes. Each session consisted of 2 blocks of 80 trials each for the behavioral study (Exp.2 and 3), 2 blocks of 60 trials each for the eye-tracking study (Exp.4 and 5), and 2 blocks of 50 trials each for the fMRI study (Exp.6). There was a short break between two blocks. The two sessions were counterbalanced across participants. It is worth noting that in Exp.2-6, one of the task aspects (loss-win or incorrect-correct) became congruent with the emphasized dimension (utility or performance) and thus became salient after emphasis. Specifically, the difference in switching frequency following loss vs. win (L-W) trials was congruent with the emphasis on utility and was thus salient when the utility was emphasized. Similarly, the difference in switching frequency following incorrect vs. correct (I-C) trials was congruent with the emphasis on performance and was thus salient when performance was emphasized.

### Subjective rating

After the Eye-tracking and fMRI experiment, participants were debriefed and required to indicate how satisfied and surprised they felt for the 8 examples of outcomes (WL, WI, LC, and LI for each session) using an 11-point analog Likert scale (0 = not at all, 10 = very intensely).

### Eye tracking apparatus and acquisition

Participants’ eye movements were recorded with a head-supported noninvasive infrared EyeLink 1000 System (SR Research). The stimuli were presented using E-prime with the EyeLink Toolbox. One eye was tracked at 1000 Hz. The participants were seated 60 cm from a computer screen in a dimly lit, sound-attenuated room. The experiment was administered on a 20-inch (40 × 30-cm screen size) Lenovo CRT display (1024 × 768 screen resolution). The eye tracker was calibrated with the built-in 9-point grid method at the beginning of the experiment. Fixation and saccades were measured using software supplied with the EyeLink eye-tracking system. Saccade detection required a deflection of > 0.1°, with a minimum velocity of 30°/s and a minimum acceleration of 8000°/s2. Fixations were defined as periods without saccades, and the objects fixated on were determined using the EyeLink event parser. The measurements were very high resolution, with resolutions as small as 5 microns (0.005 mm). All the data was exported for further analysis using Matlab 2022 (www.mathworks.com).

### Eye tracking data analysis: ROIs selection and data extraction

The eye movements were recorded continuously from the onset of the feedback delivered. Two regions of interest (ROIs) corresponding to the two cards were defined and the fixation-related eye movements landed in the two ROIs were quantitatively measured and the saccadic trajectories between them were tracked. Each ROI was a rectangle with a fixed width corresponding to the size of each card. The fixation properties of the eye movements within the ROIs were calculated for each card over the entire 1.5 s duration. Here, we mainly focused on the fixation duration (i.e., the dwell time that each subject fixates on the ROIs in each trial) and relative fixation difference between chosen and unchosen values when the feedback was delivered, and further how the fixation difference could be modulated by our salience manipulation.

### Statistics

A salience (utility vs. performance) × utility (win vs. loss trials) × performance (correct vs. incorrect trials) repeated-measures ANOVA was performed. The dependent variables were switching frequency, fixation duration, or subjective ratings. The Mauchly test was used to assess the validity of the sphericity assumption in ANOVAs. Greenhouse-Geisser corrections were used when sphericity was violated. Because we found no significant interactions between utility X performance, four types of trials (WC, WI, LC, LI) were grouped into L-W or I-C in our analysis.

### Functional magnetic resonance imaging (fMRI): data acquisition and preprocessing

MRI scanning was conducted on a 3-Tesla Tim Trio Magnetic Resonance Imaging scanner (Siemens, Germany) using a standard 12-channel head-coil system. Whole-brain data were acquired with echo-planar T2*-weighted imaging (EPI), sensitive to blood oxygenation level-dependent (BOLD) signal contrast (32 oblique axial slice; 3.9 mm thickness; 3 mm in-plane resolution; repetition time (TR) = 2150 ms; echo time (TE) = 30 ms; flip angle = 90°; FOV = 112 mm; voxel size: 3.9 × 3 × 3 mm^3^). The imaging data were acquired at a 30° angle from the anterior commissure-posterior commissure (AC-PC) line to maximize the orbital sensitivity (Deichmann, Gottfried, Hutton, & Turner, 2003). T1-weighted structural images were acquired at a resolution of 1×1×1 mm.

Neuroimaging data were preprocessed using SPM12 (www.fil.ion.ucl.ac.uk/spm/). The first five volumes were discarded to allow the MR signal to reach steady-state equilibrium. EPI images were sinc interpolated in time to correct for slice-timing differences and realigned to the first scan by rigid-body transformations to correct for head movements. Utilizing linear and nonlinear transformations and smoothing with a Gaussian kernel of full-width-half maximum 6 mm, EPI and structural images were co-registered and normalized to the T1 MNI 152 template (Montreal Neurological Institute, International Consortium for Brain Mapping). Global changes were removed by high-pass temporal filtering with a cutoff of 128s to remove low-frequency drifts in the signal.

### fMRI: imaging analysis

First, to investigate the neural activities related to outcome evaluation, we used an event-related design and constructed a general linear model (GLM) at the onset of outcome evaluation with a factorial design (win vs. loss × correct vs. incorrect) separately for each salience emphasis. Six head-motion parameters defined by the realignment were added to the model as regressors of no interest. For every voxel, multiple linear regressions were then used to generate parameter estimates for each regressor. The second-level group analysis applied a random-effects statistical model (Penny & Holmes, 2007) on the contrast images (i.e., the difference in beta values for win-loss or correct-incorrect) from the first-level analysis (two-tailed one-sample t-test). For whole-brain analysis, activations were reported if they survived P < 0.001 uncorrected, cluster size k > 20.

To further investigate how the striatum encoded outcomes and how such neural encoding was modulated by salience emphasis (M. Delgado, Locke, Stenger, & Fiez, 2003; Lim et al., 2011; J. O’Doherty et al., 2004; Oyama et al., 2015), we conducted a region-of-interest (ROI) analysis. Three ROIs including caudate, putamen, and Nucleus Accumbens (NAcc)) were obtained from the WFU PickAtlas (http://fmri.wfubmc.edu/software/PickAtlas). For ROI analysis, activations were reported if they survived P < 0.05 family-wise error (FWE) after small volume correction (SVC) at the voxel level.

Furthermore, we repeated our analyses using the actual values of the chosen and unchosen cards to examine whether there was a parametric effect of outcome differences on neural activity. It is worth noting that the value of the chosen card (i.e., ‘chosen value’ in **Fig. S5** and **Table S1**) parametrically reflected win-loss information, whereas the difference in values between the chosen and unchosen cards (i.e., ‘chosen-unchosen’ in Supplementary **Fig. S5** and **Table S1**) parametrically reflected correct-incorrect information.

Second, to investigate the neural activities related to behavioral adjustment, we used an event-related design and constructed another GLM at the onset of outcome evaluation with a factorial design (win vs. loss X correct vs. incorrect X switch vs. stay) separately for each emphasis. Eight conditions were included in the GLM as regressors depending on the category of outcome (win, loss, correct, incorrect) and subsequent behavioral choice (switch or stay): switch win, switch loss, switch correct, switch incorrect, stay win, stay loss, stay correct, and stay incorrect. Six headmotion parameters defined by the realignment were added to the model as regressors of no interest. Notably, to study the effect of salience modulation, in the first-level analysis, we used the contrast of [Switch(L-W) - Stay(L-W)] and [Switch(I-C) - Stay(I-C)] for both salience emphases, which could reveal whether salience-emphasized task dimension (i.e., [Switch(L-W) - Stay(L-W)] congruent with the emphasis on utility and [Switch(I-C) - Stay(I-C)] congruent with the emphasis on performance) could elicit a stronger neural response following behavioral switching.

Activations were reported if they survived P < 0.001 uncorrected, cluster size *k* > 20 at the whole brain level, or if they survived P < 0.05 FWE at the peak level after SVC based on the following ROIs: putamen, caudate, NAcc, and vmPFC, that were mainly reported in (Huang et al., 2020; Zink et al., 2006; Zink et al., 2004; Zink et al., 2003).

Lastly, we conducted psychophysiological interaction (PPI) analysis. Besides regional effects, the physiological connectivity between two brain regions could also vary with the psychological context, known as the psychophysiological interaction (PPI) (Friston et al., 1997). Here, we focused on the functional connectivity that was modulated by behavioral adjustment under salience modulation. We placed the seed in the vmPFC and used the contrast [Switch(L-W) - Stay(L-W)] for emphasis on utility and [Switch(I-C) - Stay(I-C)] for emphasis on performance to identify brain regions that showed differential connectivity. The first GLM was then performed with three regressors (1) the main effect of vmPFC activity (estimated volume of interest signals from a 6-mm-radius sphere), (2) the main effect of the behavioral switching effect, and (3) the interaction effect between the vmPFC and the switching effect (PPI.ppi). The interaction contrast (PPI.ppi) images of the first-level analysis were entered into one-sample t-tests for the second-level group analysis conducted with a random-effect statistical model.

## Results

### Behavior: salience emphasis modulated behavioral switching

To investigate the impact of feedback and salience emphasis on participants’ decision-making strategies, we analyzed the switching frequency of cards, i.e., choosing a different card in the next trial. The switching frequency can index the adjustment of behavior. Previous studies using reinforcement learning have consistently indicated a behavioral tendency of choosing alternative choices following loss or less optimal decisions (Cavanagh et al., 2010; Cohen & Ranganath, 2007). We here further tested whether switching frequency could be modulated by salience emphasis.

We first established the baseline performance in Exp. 1 among 21 subjects (**Fig. 2A**), where we did not have any salience emphasis. Participants switched more frequently following incorrect trials than correct trials (two-tailed paired t-test: t (20) = 4.24, P < 0.001, Cohen’s d = 0.94). However, there was no significant difference between loss trials and win trials (t (20) = 1.49, P = 0.14, d = 0.33).

**Fig. 2.**
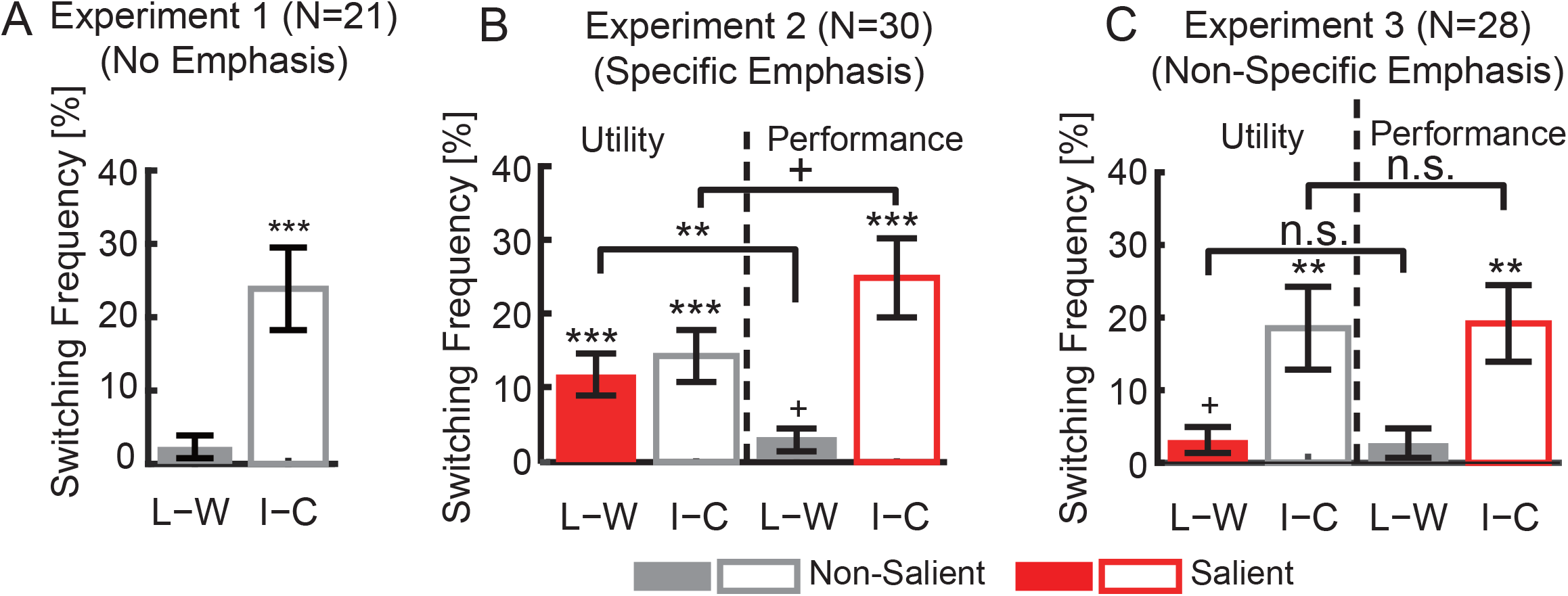
Behavioral results for Exp.1 to 3. **(A)** Exp.1. When there was no salience emphasis, participants switched more frequently following incorrect trials than correct trials but there was no significant difference between loss and win trials. The y-axis shows the percentage of trials that participants switched choice in the next trial. **(B)** Exp.2. Specific salience emphasis increased the difference in switching frequency congruent to the emphasized dimension. **(C)** Exp.3. Nonspecific salience emphasis did not increase the difference in switching frequency congruent to the emphasized dimension. Error bars denote one SEM across participants. Asterisk indicates a significant difference using two-tailed one-sample t-test: +: P < 0.1, *: P < 0.05, **: P < 0.01, and ***: P < 0.001. n.s.: not significant. Red: congruent / salient. Gray: incongruent / non-salient. Solid bars denote loss-win whereas open bars denote incorrect-correct.

We next included salience modulation in Exp.2 (**Fig. 2B**). The difference in switching frequency following loss vs. win (L-W) trials was congruent with the emphasis on utility and was thus salient when the utility was emphasized. This was confirmed by a significant interaction between salience emphasis and utility (F (1, 29) = 9.11, P = 0.0053, η_p_^2^ = 0.24). Indeed, participants switched more frequently following loss trials (mean ± SD: 53.57%±15.53%) than win trials (41.52%±17.38%) when utility was emphasized (t (29) = 4.17, P = 2.52×10^−4^, d = 0.77; see Supplementary **Fig. S1A, B** for absolute switching frequency), which was significantly stronger than the L-W effect in the no emphasis condition. However, this was not the case when performance was emphasized (t (29) = 1.94, P = 0.061, d = 0.36), suggesting that emphasis on utility specifically increased switching frequency following loss trials.

On the other hand, the difference in switching frequency following incorrect vs. correct (I-C) trials was congruent with the emphasis on performance and was thus salient when performance was emphasized. An interaction between salience and performance was also observed (F(1, 29) = 4.10, P = 0.05, η_p_^2^ = 0.13): although participants switched more frequently following incorrect trials than correct trials when either utility (25.36%±29.81%; t (29) = 4.07, P = 3.31×10^−4^, d = 0.76) or performance (14.60%±19.64%; t (29) = 4.66, P = 6.52×10^−5^, d = 0.87) was emphasized, the difference was greater when performance was emphasized (t (29) = −2.03, P = 0.05, d = −0.38), which was significantly stronger than the I-C effect in the no emphasis condition. Together, our results suggest that salience emphasis can increase the difference for the congruent (thus salient) task aspect. Therefore, adjustment of behavior (shown in switching frequency) can be modulated by salience emphasis.

In addition, we found that the switching frequency was similar for the first half vs. second half of the trials (four-way repeated-measure ANOVA of salience X utility X performance X group (first vs. second): main effect of group and interactions with the group: all Ps > 0.05), indicating no significant learning that would, in turn, modulate behavioral strategies.

Lastly, we analyzed response times (RT) for behavioral adjustment. No significant difference in RT was found when participants made either stay or switch choices (all Ps > 0.05), indicating an equal response effort that was not influenced by salience or outcome.

### Behavior: non-specific salience emphasis did not modulate behavioral switching

To test whether salience emphasis had to be specific about the chosen outcome, we conducted Exp.3 with non-specific salience emphasis—only the dimension of the emphasis (utility or performance) was indicated to participants, but not the trial-by-trial specific emphasis on the chosen outcome. Here, we found that salience modulation was abolished (**Fig. 2C**): we found no significant interaction between salience and utility (F (1, 27) = 0.039, P = 0.844, η_p_^2^ = 0.001; see Supplementary **Fig. S1C, D** for absolute switching frequency). Specifically, we found no significant difference in switching frequency between loss and win trials when either utility or performance was emphasized (both Ps > 0.1), similar to Exp.1 (**Fig. 2A**). On the other hand, no significant interaction between salience and performance was found (F (1, 27) = 0.035, P = 0.852, η_p_^2^ = 0.001, although participants switched more frequently after incorrect trials than correct trials when either utility (19.82%±31.72%) or performance (21.39%±29.75%) was emphasized. Therefore, when salience emphasis was not specific to the outcome, there was no significant difference between different salience emphases. Together, our results suggest that non-specific salience emphasis did not modulate behavioral switching.

### Eye-movement results: salience enhanced the fixation difference along the salient dimension

We first replicated our behavioral result in Exp.2 with an independent sample of 48 eye-tracking participants in Exp.4. Specifically, we identified a significant interaction between salience and utility (F (1, 47) = 11.33, P = 0.002, η_p_^2^ = 0.90, **Fig. 3A1**). When utility was emphasized, participants switched more frequently following loss trials than win trials (t (47) = 3.83, P < 0.001, d = 0.55; see Supplementary **Fig. S1E, F** for absolute switching frequency), but not when performance was emphasized (t (47) = 0.45, P = 0.65, d = 0.06). The difference between the two emphases was significant: t (47) = 3.36, P = 0.001, η_p_^2^ = 0.49.

**Fig. 3.**
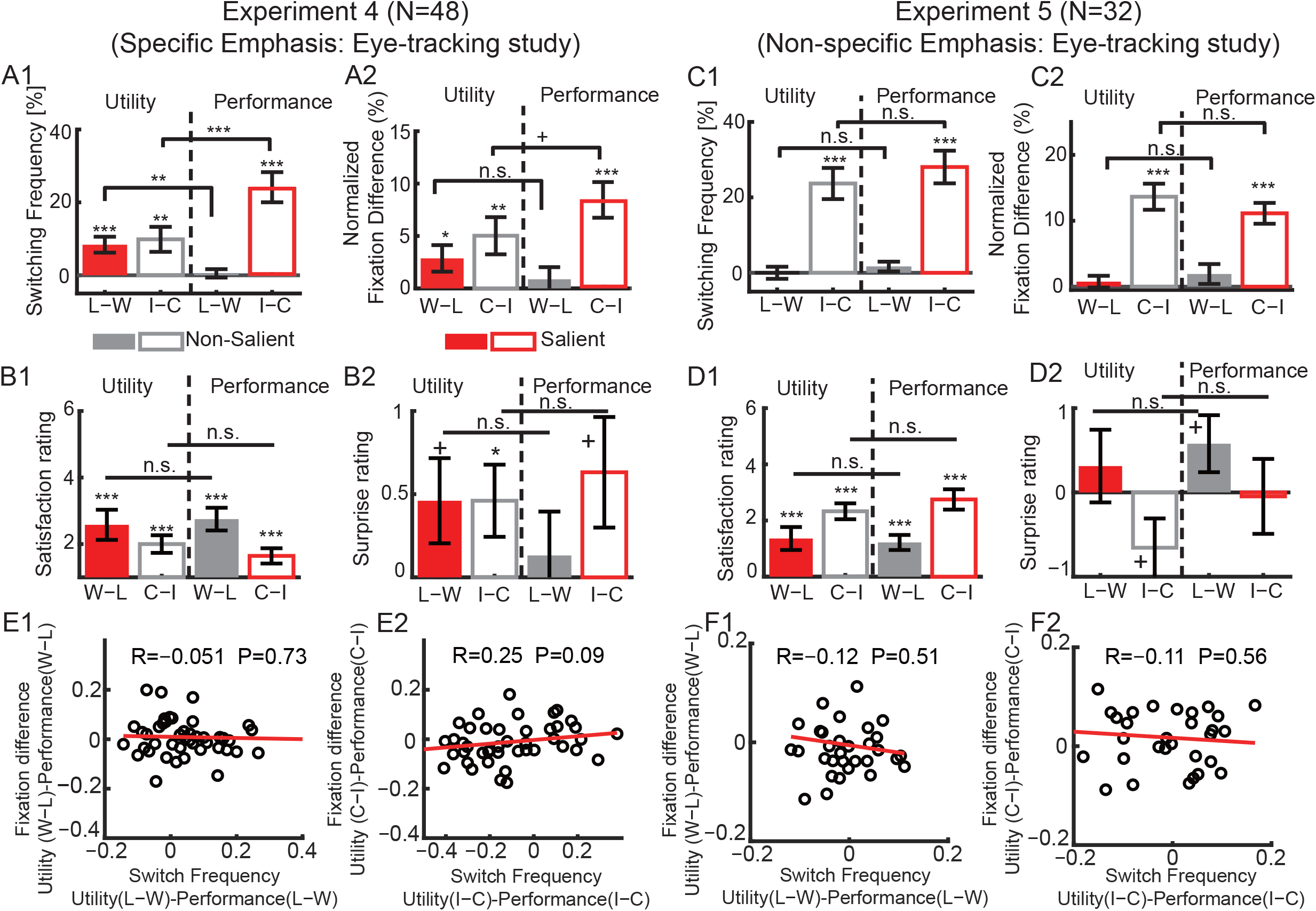
Behavioral and eye-tracking results for Exp.4 and 5. **(A1)** Specific salience emphasis increased the difference in switching frequency congruent to the emphasized dimension. **(A2)** Specific salience emphasis increased the fixation difference between chosen and nonchosen values congruent to the emphasized dimension. **(B1)** Specific salience didn’t change satisfaction ratings. **(B2)** Specific salience didn’t change surprise ratings. **(C1)** Non-specific salience emphasis did not increase the difference in switching frequency congruent to the emphasized dimension. **(C2)** Nonspecific salience emphasis did not increase the fixation difference between chosen and unchosen values congruent to the emphasized dimension. **(D1)** Non-specific salience didn’t change satisfaction ratings along the emphasized dimension. **(D2)** Non-specific salience didn’t change surprise ratings along the emphasized dimension. **(E1-E2)** Specific emphasis. **(E1)** The correlation between salience-guided utility-specific fixation difference and salience-modulated utility-specific behavioral switching didn’t reach significance. **(E2)** The correlation between salience-guided performance-specific fixation difference and salience-modulated performance-specific behavioral switching was marginally significant. **(F1-F2)** Non-specific emphasis. **(F1)** No significant correlation was identified between non-specific utility dimension-guided fixation difference and non-specific utility dimension-based behavioral switching. **(F2)** No significant correlation was identified between non-specific performance dimension-guided fixation difference and non-specific performance dimension-based behavioral switching. Error bars denote one SEM across participants. Asterisk indicates a significant difference using two-tailed one-sample t-test: +: P < 0.1, *: P < 0.05, **: P < 0.01, and ***: P < 0.001. n.s.: not significant. Red: congruent / salient. Gray: incongruent / non-salient. Solid bars denote loss-win whereas open bars denote incorrect-correct.

In addition, we identified a significant interaction between salience and performance (F (1, 47) = 16.67, P < 0.001, η_p_^2^ = 0.98, **Fig. 3A1**). Participants switched more frequently following incorrect than correct trials when either utility (t (47) = 2.87, P = 0.007, d = 0.41) or performance (t (47) = 5.84, P < 0.001, d = 0.85) was emphasized. The difference was enhanced along the salient dimension: t (47) = −4.08, P < 0.001, d = −0.59).

Notably, we observed qualitatively similar results between Exp.2 (Fig.3A1) and 4 (Fig.2B) (twotailed two-sample t-test; all Ps > 0.2). Therefore, we confirmed our results with an independent population of participants that behavioral switching can be modulated by salience emphasis.

We then investigated the fixation difference between chosen and unchosen cards to quantify the attentional deployment during outcome evaluation. We first checked the total fixation duration during outcome evaluation and found a generally similar duration of fixation for different types of outcomes (see Supplementary **Fig. S2A1, B1**). A divisive normalization procedure was performed to get the normalized fixation difference between two cards, in which the relative fixation differences (chosen - unchosen) were divided by the total fixation of two cards.

A qualitatively similar pattern of fixation deployment (**Fig. 3A2**) was observed as the behavioral switching (**Fig. 3A1**). Specifically, the normalized fixation difference was enhanced after trials with win feedback compared to loss when utility was emphasized (t (47) = 2.26, P = 0.028, d = 0.33; see Supplementary **Fig. S2A2** and **A3** for absolute changes), but not when performance was emphasized (t (47) = 0.75, P = 0.55, d = 0.10, see Supplementary **Fig. S2B2** and **B3** for absolute changes). However, the difference between the two emphasizes didn’t reach significance: t (47) = 1.11, P = 0.27, d = 0.16), which was also indicated by the non-significant interaction between salience and utility (F (1, 47) = 1.23, P = 0.27, η_p_^2^ = 0.19, **Fig. 3A2**).

Moreover, the normalized fixation difference was enhanced after trials with correct feedback compared to incorrect when either utility (t (47) = 2.83, P = 0.007, d = 0.41) or performance (t (47) = 5.84, P < 0.001, d = 0.85) was emphasized. A marginally significant difference between the two emphasizes was observed: t (47) = −1.71, P = 0.09, d = −0.25), which was also indicated by a weak interaction between salience and performance (F (1, 47) = 2.95, P = 0.09, η_p_^2^ = 0.39, **Fig. 3A2**).

In addition to behavioral switching, we also investigated the subjective pleasantness and surprise ratings for each experimental condition in Exp.4. As expected, participants were more satisfied in win trials than in loss trials, and more satisfied in correct trials than in incorrect trials, when either utility or performance was emphasized (**Fig. 3B1**; three-way repeated-measure ANOVA of salience X utility X performance: main effect of utility: F(1, 37) = 54.01, P < 0.001, η_p_^2^ = 1.0; main effect of performance: F(1, 37) = 81.04, P < 0.001, η_p_^2^ = 1.0). However, no significant interaction effects between salience and utility or performance were observed (all Ps > 0.2). For the self-reported surprise, participants were more satisfied in correct trials than in incorrect trials (**Fig. 3B2**; three-way repeated-measure ANOVA of salience X utility X performance: main effect of performance: F (1, 37) = 5.41, P = 0.026, η_p_^2^ = 0.62). No significant main effect of utility (F (1, 37) = 2.49, P = 0.12, η_p_^2^ = 0.33) or interactions (**Fig. 3B2**; all Ps > 0.1) were found. Therefore, the salience-guided behavioral adjustments cannot be simply attributed to the difference in subjective feelings towards outcomes.

Lastly, we analyzed response times (RT) for behavioral adjustment. No significant difference in RT was found when participants made switch choices under two emphases (see Supplementary **Fig. S3A1-A3;** all Ps > 0.1). Interestingly, participants tended to be slower following incorrect feedback than correct feedback when made a stay choice under performance emphasis but tended to be faster under utility emphasis, yielding a significant interaction between salience and performance (F (1,47) = 5.52, P = 0.023, η_p_^2^ = 0.63).

### Eye-movement results: non-specific salience didn’t modulate the fixation deployment

Similarly, we replicated our behavioral findings in Exp.3 using an independent group of 32 eye-tracking subjects in Exp.5. We found that salience modulation was abolished when the emphasis is non-specific (**Fig. 3C1**). Specifically, no significant interaction between salience and utility (F (1, 31) = 0.58, P = 0.45, η_p_^2^ = 0.11; see Supplementary **Fig. S1G, H** for absolute switching frequency) and no significant main effect of valence (F (1, 31) = 0.70, P = 0.40, η_p_^2^ = 0.12) on switching frequency was observed. Although participants switched more frequently after incorrect trials than correct trials (main effect of performance: (F (1, 31) = 46.01, P < 0.001, η_p_^2^ = 1), no significant interaction was found between salience and performance (F (1, 31) = 1.41, P = 0.24, η_p_^2^ = 0.21). Again, our results suggest that non-specific salience emphasis did not modulate behavioral switching. Notably, we observed qualitatively similar results between Exp.5 (**Fig. 3C1**) and 3 (**Fig. 2C**) (two-tailed two-sample t-test; all Ps > 0.2).

We then investigated the fixation difference between chosen and unchosen values. A qualitatively similar pattern of fixation deployment (**Fig. 3C2**) was observed as the behavioral switching in Exp. 5 (**Fig. 3C1**). Specifically, the normalized fixation difference was unchanged between trials with win and loss feedback (main effect of utility: F (1, 31) = 2.55, P = 0.12, η_p_^2^ = 0.34; see Fig. S2C2-D3 for absolute changes). Moreover, there was no significant interaction between salience and utility (F (1, 31) = 0.40, P = 0.52, η_p_^2^ = 0.09, **Fig. 3C2**). Although the normalized fixation difference was enhanced after trials with correct feedback compared to incorrect (main effect of performance: F (1, 31) = 58.73, P < 0.001, η_p_^2^ = 1), no significant difference between two emphasizes was observed (F (1, 31) = 2.66, P = 0.11, η_p_^2^ = 0.35, **Fig. 3C2**).

Next, we investigated the subjective pleasantness and surprise ratings for each experimental condition. Similar to the findings in Exp.4, participants were more satisfied in win trials than in loss trials, and more satisfied in correct trials than in incorrect trials, when either utility or performance was emphasized (**Fig. 3D1**; three-way repeated-measure ANOVA of salience X utility X performance: main effect of utility: F(1, 31) = 17.58, P < 0.001, η_p_^2^ = 0.98; main effect of performance: F(1, 31) = 69.03, P < 0.001, η_p_^2^ = 1.0). However, no significant interaction effects between salience and utility or performance were observed (all Ps > 0.1). For the self-reported surprise, no significant main effects or interactions were found (**Fig. 3B2**; all Ps > 0.1). Overall, we have observed similar results in subjective feelings regardless of the specification of outcomes that are also independent of behavioral switching.

Lastly, we analyzed response times (RT) for behavioral adjustment. No significant difference in RT was found when participants made either switch (see Supplementary **Fig. S3C1-C3;** all Ps > 0.47) or stay (**Fig. S3D1-D3;** all Ps > 0.43) choices, indicating an equal response effort that was not influenced by salience or outcome.

### Eye-movement results: behavioral switching was correlated with attention deployment

We have identified similar patterns of behavioral switching and attentional deployment. To examine the role of attention in salience effects, we first performed a Pearson correlation analysis between the behavioral switching and attentional deployment for each condition.

When emphasizing utility, the switching difference along the salient utility dimension (L-W) was positively correlated with the normalized fixation difference (Chosen-Unchosen) under the same dimension (W-L) (see Supplementary **Fig. S4A1;** r = 0.29, P=0.046). Moreover, the switching difference along the non-salient performance dimension (I-C) was positively correlated with the normalized fixation difference (Chosen-Unchosen) under the same dimension (C-I) (see Supplementary **Fig. S4A2;** r = 0.4, P=0.005).

When emphasizing performance, the switching difference along the salient performance dimension (I-C) was positively correlated with the normalized fixation difference (Chosen-Unchosen) under the same dimension (C-I) (see Supplementary **Fig. S4A4;** r =0.33, P=0.023). No correlation was identified between the switching difference along the non-salient utility dimension (L-W) and the normalized fixation difference (Chosen-Unchosen) under the same dimension (W-L) (see Supplementary **Fig. S4A3;** r = −0.19, P=0.2). Altogether, our results tend to support that specific feedback modulated attention deployment and further guided behavioral adjustments.

Notably, no direct correlation was observed between salient utility-modulated fixation difference [Utility (W-L) – Performance (W-L)] and salient utility-modulated behavioral switching [Utility (L-W) – Performance (L-W)] (**Fig. 3E1**; r =-0.05, P=0.73). Moreover, only a marginally significant correlation was observed for salient performance-modulated fixation difference [Utility (C-I) – Performance (C-I)] and salient performance-modulated behavioral switching [Utility (I-C) – Performance (I-C)] (**Fig. 3E2**; r =0.25, P=0.09). These results may indicate that the observed salience-guided fixation deployment didn’t linearly guide the salience-based behavioral switching.

Moreover, no significant correlation between switching difference and attentional deployment was identified under the non-specific salience emphasis (**Fig. 3F1-F2:** all P values > 0.5; Supplementary **Fig. S4B1-B4**, all P values > 0.2).

Lastly, a general linear mixed model was used to predict the behavioral strategy using chosen value, unchosen value, and normalized fixation difference as independent variables for each subject. We have identified a significant role of chosen value, unchosen value, and normalized fixation difference on the behavioral strategy when either utility (see Supplementary, **Fig. S4C1**) or performance (**Fig. S4C2**) was emphasized. However, no significant effect of normalized fixation difference was identified when the emphasis was non-specific (see Supplementary **Fig. S4D1, D2**).

### Replication in the fMRI study

Notably, we further replicated the salience effect using color emphasis by an independent sample of 25 fMRI participants (Exp.6). Qualitatively the same results were observed (**Fig. 4A**): when utility was emphasized, participants switched more frequently following loss trials (52.85% ±12.16%) than win trials (44.35%±16.73%; t (24) = 2.76, P = 0.011, d = 0.56; see Supplementary **Fig. S1I, J** for absolute switching frequency), but not when performance was emphasized (t (24) = 0.30, P = 0.76, d = 0.06; interaction: F(1, 24) = 5.12, P = 0.033, η_p_^2^ = 0.58). In addition, participants switched more frequently following incorrect than correct trials when either utility (9.97%±22.53%; t (24) = 2.21, P = 0.03, d = 0.45) or performance (16.12%±24.92%; t (24) = 3.23, P = 0.004, d = 0.66) was emphasized (the difference between two emphases: t (24) = −1.60, P = 0.12, d = −0.33). Notably, we observed qualitatively similar results between Exp.2 and 6 for each condition (two-tailed two-sample t-test; all Ps > 0.2).

**Fig. 4.**
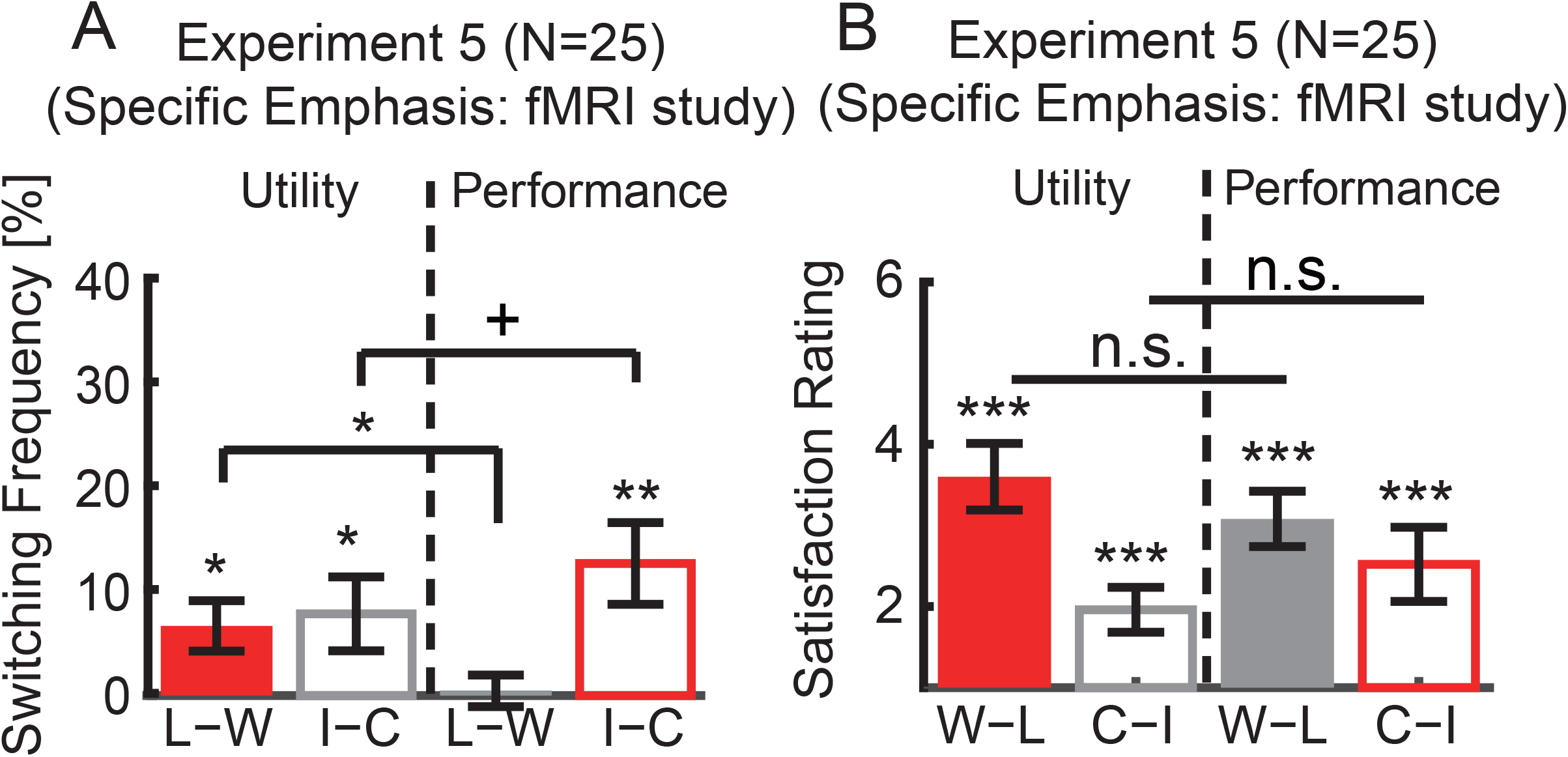
Behavioral results for fMRI study. **(A)** Exp.6. (fMRI participants) replicated the results from Exp.2 (behavioral participants). **(B)** Satisfaction rating from fMRI participants. Error bars denote one SEM across participants. Asterisk indicates a significant difference using two-tailed one-sample t-test: +: P < 0.1, *: P < 0.05, **: P < 0.01, and ***: P < 0.001. n.s.: not significant. Red: congruent / salient. Gray: incongruent / non-salient. Solid bars denote loss-win whereas open bars denote incorrect-correct.

Similarly, we investigated subjective pleasantness and surprise ratings of outcomes in fMRI participants. As expected, participants were more satisfied in win trials than in loss trials, and more satisfied in correct trials than in incorrect trials, when either utility or performance was emphasized (**Fig. 4B**; three-way repeated-measure ANOVA of salience X utility X performance: main effect of utility: F(1, 24) = 116.18, P = 1.11×10^−10^, η_p_^2^ = 1.0; main effect of performance: F(1, 24) = 49.30, P = 2.92×10^−7^, η_p_^2^ = 1.0). However, no significant interactions between salience and utility or performance were observed (all Ps > 0.1). For the self-reported surprise, no significant main effects or interactions were found (all Ps > 0.1). Therefore, the salience-guided behavioral adjustments cannot be simply attributed to the difference in subjective feelings towards outcomes.

### fMRI: the striatum subregions encoded salience-modulated outcome evaluation

We next investigated the neural substrates underlying this behavior. The striatum showed a greater activity associated with win outcome vs. loss outcome when either utility (**Fig. 5A**) or performance (**Fig. 5B**) was emphasized (see **Table 1** and Supplementary **Table S1** for full statistics). The difference between win vs. loss outcomes became greater when the utility was emphasized (congruent and salient) compared with when performance (incongruent and non-salient) was emphasized, suggesting that the congruent and thus salient task aspect could enhance the striatum’s coding of utility. To test the statistical significance of the salience effect on neural activity, we conducted an ROI analysis (see **Methods** for choice of ROIs) in the striatum subregions. We found that the dorsal striatum (particularly of the **left caudate**) had a greater activity for win-loss when the utility was emphasized than when performance was emphasized (**Fig. 5E**; peak: Montreal Neurological Institute (MNI) coordinate: x = −12, y = 18, z = 3, 10 voxels, FWE P < 0.05, small volume corrected (SVC); **Fig. 5G**; two-tailed one-sample t-test against 0: utility: t (24) = 4.50, P = 1.45×10^−4^, d = 0.91; performance: t (24) = 2.87, P = 0.008, d = 0.58; two-tailed paired t-test between two emphases: t (24) = 1.80, P = 0.08, d = 0.37; see also **Table 1**), suggesting the left caudate selectively encodes salience-modulated utility information.

**Fig. 5.**
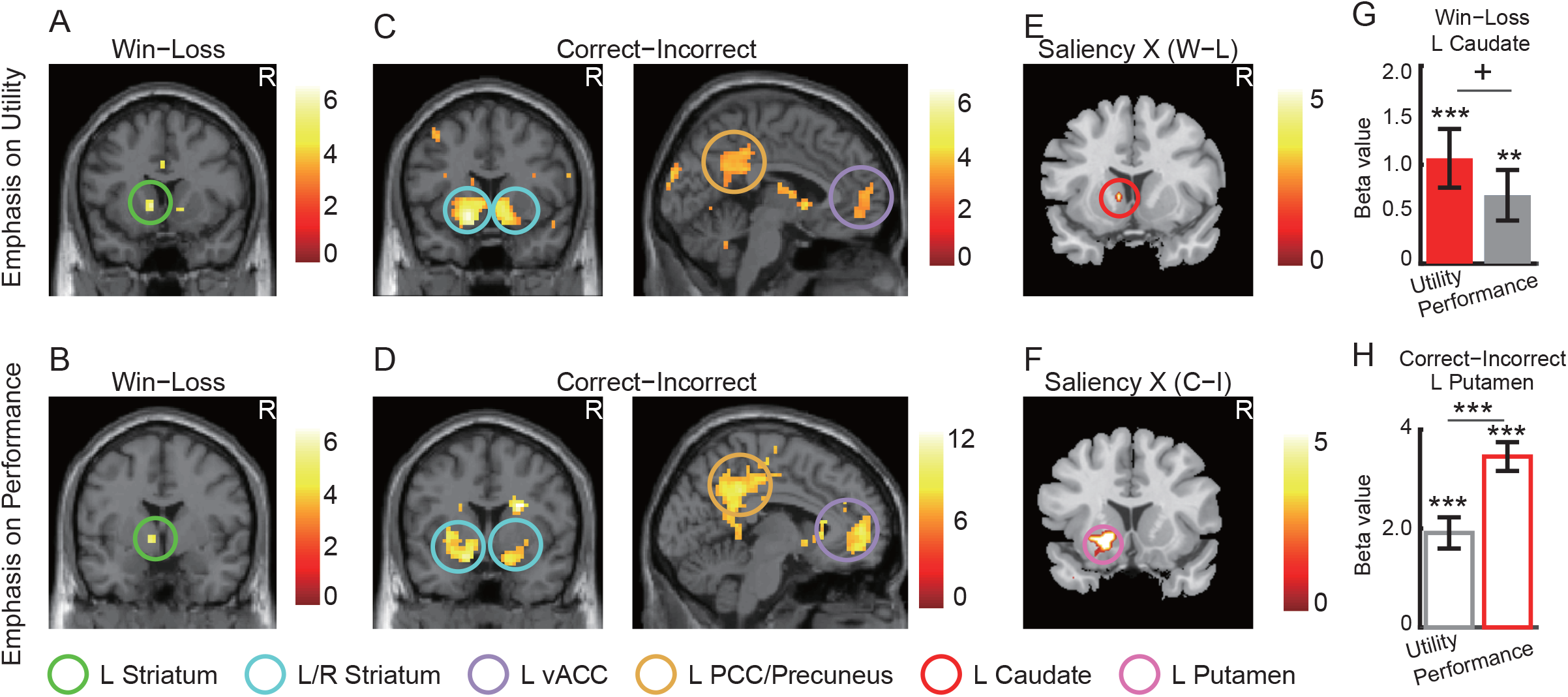
Salience-modulated outcome evaluation. **(A)** The left caudate (x = −9, y = 12, z = 0) encoded utility (win-loss) when utility was emphasized. **(B)** The left striatum (x = −9, y = −3, z = −3) also encoded utility (win-loss) when performance was emphasized. **(C, D)** Both the left and right striatum encoded performance (correct-incorrect) when either utility **(C)** or performance **(D)** was emphasized. **(E)** The left caudate (x = −12, y = 18, z = −3) encoded the interaction between salience and utility (win-loss). **(F)** The left putamen (x = −18, y = 12, z = −6) encoded the interaction between salience and performance (correct-incorrect). **(G, H)** Parameter estimate (beta values). The left bar shows parameter estimates for emphasis on utility and the right bar shows parameter estimates for emphasis on performance. The bars show the average beta values of all voxels from the ROI. Red: congruent / salient. Gray: incongruent / non-salient. Solid bars denote win-loss whereas open bars denote correct-incorrect. The generated statistical parametric map was superimposed on anatomical sections of the standardized MNI T1-weighted brain template. Images are in neurological format with participants left on the image left. L: left, R: right. Activations were shown at P < 0.001 uncorrected. Error bars denote one SEM across participants. Asterisk indicates a significant difference using a two-tailed one-sample t-test: **: P < 0.01. ***: P < 0.001.

**Table 1.** Brain areas are modulated by salience-modulated outcome evaluation and behavioral adjustment. All values are P < 0.001 uncorrected at the peak voxel level. * indicates P < 0.05 and ** indicates P < 0.01 family-wise error (FWE) after small volume correction (SVC).

|  | Contrast | Brain Region | Z-score | Peak Coordinate<br>MNI (X Y Z) |  |  | Volume<br>(voxel) |
| --- | --- | --- | --- | --- | --- | --- | --- |
| Salience-modulated outcome evaluation |  |  |  |  |  |  |  |
| Salience X<br>Utility | Utility (W–L)<br>–<br>Performance<br>(W–L) | L Caudate head* | 2.98* | –12 | 18 | –3 | 2 |
| Salience X<br>Performance | Utility (C–I)<br>–<br>Performance<br>(C–I) | L Caudate*<br>L Putamen*<br>R Putamen* | 3.40*<br>3.43*<br>3.39* | –18<br>–18<br>27 | 21<br>12<br>0 | 9<br>–6<br>–6 | 3<br>5<br>5 |
| Salience-modulated behavioral adjustment |  |  |  |  |  |  |  |
| Salience X<br>Utility X<br>Strategy | Utility<br>(Switch –Stay) X (L–W)<br>–<br>Performance<br>(Switch –Stay) X (L–W) | L vACC* | 3.52* | –15 | 42 | 0 | 5 |
| Salience X<br>Performance X<br>Strategy | Utility<br>(Switch –Stay) X (I–C)<br>–<br>Performance<br>(Switch –Stay) X (I–C) | R vmPFC* | 3.84* | 3 | 54 | 0 | 22 |
| “High order” PPI Results |  |  |  |  |  |  |  |
| Salience X<br>Utility X<br>Strategy | Utility<br>(Switch –Stay) X (L–W)<br>–<br>Performance<br>(Switch –Stay) X (L–W) | L Nucleus<br>Accumbens** | 3.28** | –15 | 3 | –15 | 5 |
| Salience X<br>Performance X<br>Strategy | Utility<br>(Switch –Stay) X (I–C)<br>–<br>Performance<br>(Switch –Stay) X (I–C) | R dmPFC/Middle<br>Frontal Gyrus** | 4.55** | 15 | 42 | 27 | 12 |

Similarly, the striatum also showed a greater activity associated with correct outcome vs. incorrect outcome, when either utility (**Fig. 5C**) or performance (**Fig. 5D**) was emphasized (see **Table 1** and Supplementary **Table S1** for full statistics). Other brain regions that were activated included the ventral medial prefrontal cortex (vmPFC). However, the difference between correct vs. incorrect outcomes became greater in the striatum when performance was emphasized that involved both the dorsal (left **caudate**) and ventral (bilateral **putamen**) striatum, suggesting that the congruent and thus salient task aspect could enhance the coding of performance as well as represented by both ventral and dorsal striatum. ROI analysis further confirmed the results and revealed a significant difference between salience emphases (the **left putamen**; **Fig. 5F**; peak: x = −18, y = 12, z = −6, Z = 3.68, 27 voxels, FWE P < 0.05, SVC; **Fig. 5H**; two-tailed paired t-test between two emphases: t (24) = 4.51, P = 1.41×10^−4^, d = 0.92; see also **Table 1**). Therefore, both ventral and dorsal striatum represent salience-modulated performance information.

It is worth noting that no significant correlation was observed between the neural response from the striatum (left caudate or putamen) and the corresponding behavioral switching under salience manipulation. Moreover, although the above analyses were performed categorically (i.e., win, loss, correct, incorrect), we repeated our analyses using parametric effects on outcome difference (i.e., using actual payoff values) and we derived qualitatively the same results (see Supplementary **Fig. S5** and **Table S2**). Together, our results suggested that the striatum not only encoded utility and performance but also could be modulated by salience emphasis. This was in accordance with behavior (**Fig. 4**) and might explain salience-modulated behavioral adjustment, a point that we will elucidate next.

### fMRI: the vmPFC encoded salience-modulated behavioral adjustment

We next investigated the brain regions that may encode behavioral adjustment following outcome evaluation under salience modulation, which corresponded to our observed behavior (**Fig. 4**). We identified the brain regions that were activated under the contrasts of [Switch(L-W) - Stay(L-W)] and [Switch(I-C) - Stay(I-C)] for both salience manipulations. Interestingly, the vmPFC was activated during behavioral switching following both utility and performance information but showed a stronger response following information congruent with the salience manipulation (**Fig. 6A-F**). ROI analysis further confirmed the results and revealed a significant difference between salience-modulated behavioral adjustment (the vmPFC; **Fig. 6E**; peak: x = −15, y = 42, z = 0, Z = 3.52, 6 voxels, FWE P < 0.05, SVC; **Fig. 6F**; peak: x = 3, y = 54, z = 0, Z = 3.84, 22 voxels, FWE P < 0.05, SVC). Therefore, the vmPFC represents salience-driven behavioral adjustments.

**Fig. 6.**
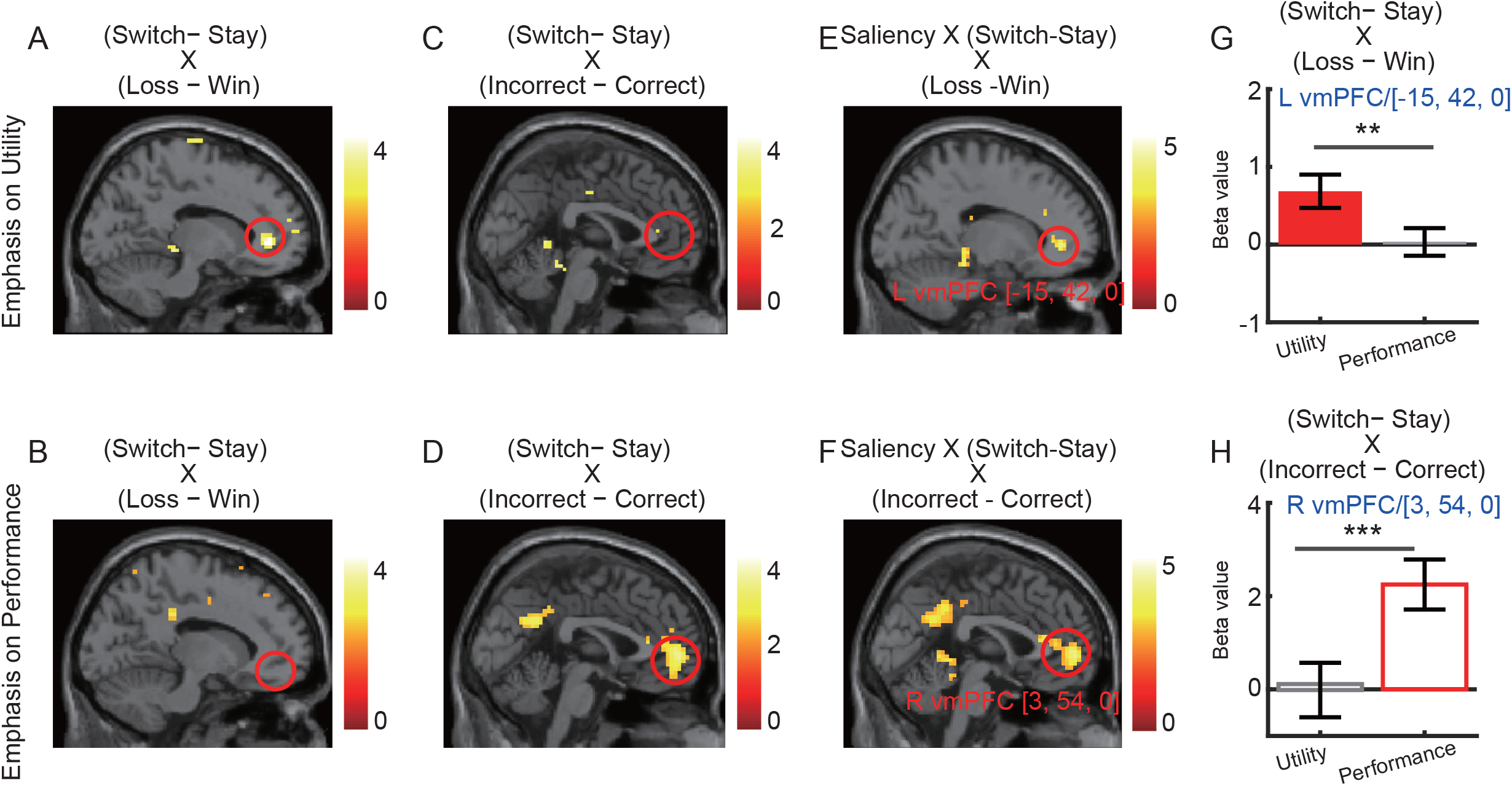
Salience-modulated behavioral adjustment. **(A, B)** When utility was emphasized, the left vmPFC was activated under behavioral adjustment following utility information [Switch(L-W) - Stay(L-W)], while no brain regions showed any significant activation when performance was emphasized. **(C, D)** When performance was emphasized, the behavioral adjustment following the performance information [Switch(I-C)-Stay(I-C)] involved the right vmPFC. **(E, F)** The interaction between salience and utility-based behavioral switching and performance-based behavioral switching. **(G, H)** Beta values in vmPFC show the interaction between salience and behavioral switching. The generated statistical parametric map was superimposed on anatomical sections of the standardized MNI T1-weighted brain template. Images are in neurological format with participants left on the image left. L: left, R: right. Activations were shown at P < 0.005 uncorrected. Asterisk indicates a significant difference: ***: P < 0.001. Each dot shows an average beta value of all voxels from the ROI.

### fMRI: behavioral adjustment modulated functional connectivity between the vmPFC and NAcc/dmPFC

Lastly, to further explore whether the vmPFC was functionally connected with other brain regions and whether such connectivity could be modulated by salience-modulated behavioral adjustment, we performed a classical PPI analysis with the vmPFC (utility: MNI peak: x=-15, =42, z=0; performance: x=3, =54, z=0) as the seed and the signals from a 6-mm-radius sphere around the seed as a volume of interest (VOI). However, no brain regions were significantly activated and showed connectivity with the vmPFC under two emphases.

A further “high-order” PPI model was constructed in which the salience-guided behavioral switching after utility or performance from each participant was put into one vector to capture brain regions that may respond to the main contrast of PPI.ppi generated from the primary PPI model. This is indeed a regression model used to identify the regions functionally connected with the vmPFC, and further, their connectivity was modulated by the behavioral switching for each salience manipulation. We found that the left vmPFC was co-activated with the left NAcc (**Fig. 7A**; MNI peak: x = −15, y = 3, z = −15, Z = 3.28, 5 voxels, FWE P < 0.01, SVC) when emphasizing on utility. A Pearson correlation analysis further confirmed a high inter-participant negative correlation between the strength of vmPFC-NAcc connectivity and the behavioral switching following utility (**Fig. 7B**). Moreover, the right vmPFC was co-activated with the right dmPFC (**Fig. 7C**; peak: x = 15, y = 42, z = 27, Z = 4.55, 12 voxels, FWE P < 0.01, SVC) when emphasizing on performance. A Pearson correlation analysis further confirmed a high intra-participant negative correlation between the strength of vmPFC-dmPFC connectivity and the behavioral switching following performance (**Fig. 7D**).

**Fig. 7.**
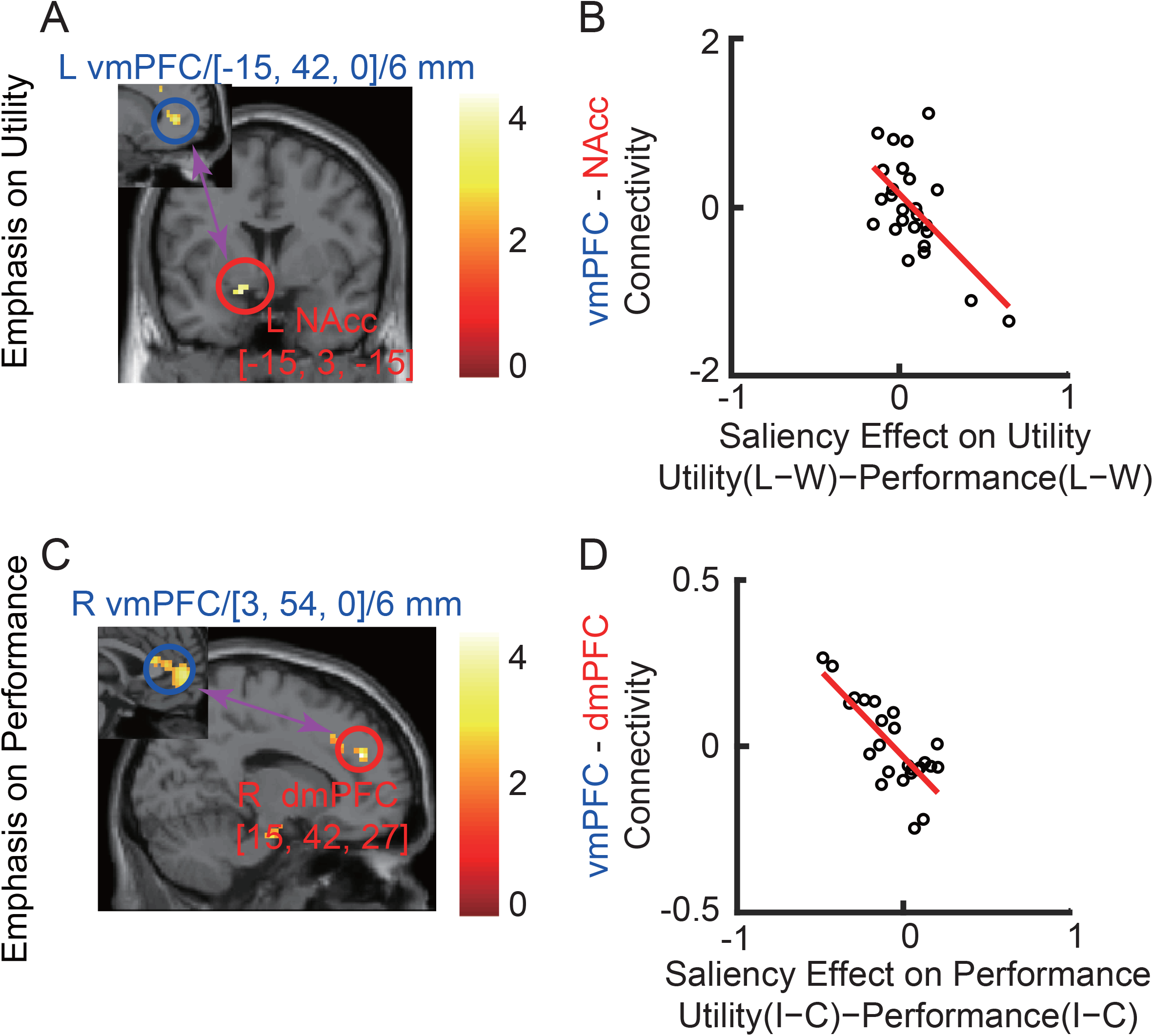
Regional activity on behavioral adjustment and high-order PPI results. **(A)** When utility was emphasized, the left vmPFC (x = −15, y = 42, z = 0) encoded behavioral adjustment following utility feedback, and its connectivity to nucleus accumbens (NAcc) (x = −15, y = 3, z = −15) was negatively modulated salience-guided behavioral adjustments. Activations were shown at P < 0.005 uncorrected. **(B)** When performance was emphasized, the right vmPFC (x = 3, y = 54, z = 0) encoded behavioral adjustment following performance feedback, and its connectivity to dmPFC (x = 15, y = 42, z = 27) was negatively modulated by the behavioral adjustments. Activations were shown at P < 0.005 uncorrected.

Together, our results indicated that the frontostriatal neural circuit could be modulated by salience-driven behavioral adjustment.

## Discussion

In this study, we investigated the cognitive and neural bases of salience-driven incidental learning. Consistent with our hypothesis, participants showed salience-modulated behavioral adjustment in experiments that had a specific association between visual salience and outcome, but not in experiments the association between visual salience and outcome was non-specific. These findings suggested that salience emphasis exerted influences through specific mapping of bottom-up visual cues with feedback, yielding feedback-specific salience-modulated behavioral adjustments. The striatum subregions encoded salience-modulated outcome evaluation, while the vmPFC encoded salience-guided behavioral adjustment. The functional connectivity of vmPFC-NAcc accounted for the inter-individual difference in utility-driven behavioral adjustment, while the functional connectivity of vmPFC-dmPFC accounted for the inter-individual difference in performance-driven behavioral adjustment. Together, our results suggest that feedback-specific visual salience modulates behavioral adjustment, presumably by adjusting how outcome information is weighted in the evaluation process, and further guides incidental learning, processes that may involve attention and the frontal-striatal valuation systems.

### *Visual* salience *and decision making*

Traditional decision-making studies typically present participants with options that are well balanced in visual salience to rule out confounding variables from low-level perception on brain activity and behavior. However, options in real life rarely appear in a visual vacuum. For example, an aircraft pilot needs to encode information from the instrument panels and make critical decisions after weighing the importance of the information. Our study mainly conveys that visual salience interplays with value-based decision-making. In line with this viewpoint, it has been demonstrated that perceptual salience competes with the expected value, and both influence the saccadic endpoint within an object (Krajbich, Armel, & Rangel, 2010; Schütz, Trommershäuser, & Gegenfurtner, 2012; Towal, Mormann, & Koch, 2013). Moreover, motivationally salient stimuli, such as items previously associated with rewards, can bias visual attention and subsequent decision strategy (Hickey, Chelazzi, & Theeuwes, 2010). A reward-associated distractor can change saccade trajectories even when participants expect this object and try to ignore it (Hickey & van Zoest, 2012).

Additionally, manipulating the relative level of visual attention between two alternative options can influence subsequent choices (Armel, Beaumel, & Rangel, 2008). Similar to our present findings, behavioral economic studies have suggested that deemphasizing a stock’s purchase price can substantially reduce stockers’ propensity to sell risky assets with capital gains (Frydman & Rangel, 2014). However, the above-mentioned salience modulation could be confounded by the simple demand characteristics effect where individuals pay more (or less) attention to the emphasized (or non-emphasized) dimension. Our results have further extended previous studies by showing that outcome salience, even when it is redundant in nature, has an impact on subsequent decisions, which is different from the effect of the demanding characteristics. We have further shown the specificity of salience modulation: only when salience emphasis is specific about the chosen outcome, it can exert an impact on subsequent choices, that are represented by the striatal and frontal subregions.

### The striatum subregions encode salience-modulated outcome evaluation

Our neuroimaging results have revealed that the striatum subregions were involved in both utility (i.e., dorsal striatum: the caudate) and performance (both dorsal and ventral striatum: bilateral caudate, putamen, and NAcc) evaluation, and their activities were further modulated by salience emphasis. The human striatum has long been implicated in value-based decision-making, and it can be significantly activated by the positive versus negative feedback (Becker, Nitsch, Miltner, & Straube, 2014). However, it has been argued that the striatum is not only engaged in reward processing but also encodes stimulus salience (M. R. Delgado, 2007; Guitart-Masip, Bunzeck, Stephan, Dolan, & Düzel, 2010; Jensen et al., 2003; Oyama et al., 2015; Zaehle et al., 2013; Zink et al., 2006; Zink et al., 2004; Zink et al., 2003). In particular, the ventral striatum is shown to be modulated by both value and visual salience (Litt, Plassmann, Shiv, & Rangel, 2011), and it encodes attention-guided relative-value signals (Lim et al., 2011), anticipated aversive stimuli (Jensen et al., 2003), salient non-rewarding stimuli (Zink et al., 2003), and salient prediction error (Metereau & Dreher, 2013). Our results have confirmed the crucial role of the striatum in salience processing.

Moreover, the dorsal striatum represents visual salience-based outcome evaluation regardless of the utility or performance information (feedback-general), while the ventral striatum only represents salience-based performance information (feedback-specific). This is in line with the studies on goal-directed and habitual processes, in which the ventral striatum is involved in both, while the dorsal striatum only encodes habitual learning (Huang et al., 2020). Our results indicate that feedback-general salience may interact with the habitual learning process to guide outcome evaluation spontaneously by recruiting the dorsal striatum. In contrast, the salience-guided performance-specific outcome evaluation is more goal-directed compared to the utility-specific outcome evaluation given that the ventral striatum is only activated for performance emphasis and represents habitual learning process as well. Based on these, our findings have advanced the neurocognitive understanding of salience processing, by showing that salience-driven valuation may rely on the specificity of feedback and interplay with both goal-directed and habitual learning processes.

Another line of evidence from human electrophysiology studies has identified two ERP components, the feedback-related negativity (FRN) and the P300, that are sensitive to the reward-based outcome evaluation (Cohen & Ranganath, 2007; Gehring & Willoughby, 2002; Nieuwenhuis, Yeung, Holroyd, Schurger, & Cohen, 2004; Yeung & Sanfey, 2004). In particular, the FRN discriminates monetary outcomes with different salience levels that are rendered by levels of perceptual noise, suggesting an interaction between stimulus salience and the value computation (Lou, Hsu, & Sajda, 2015). Source localization of the EEG components has suggested that the FRN originates from cortical regions such as the MFG that receive dopaminergic projections from the basal ganglia (including the striatum) and reflects activity in the mesocorticolimbic reward circuits (Becker et al., 2014; Carlson, Foti, Mujica-Parodi, Harmon-Jones, & Hajcak, 2011; M. Delgado et al., 2003; Knutson, Fong, Bennett, Adams, & Hommer, 2003; Nieuwenhuis et al., 2005). A simultaneous EEG-fMRI study showed that surprise-like salience signals are directly projected to the source region of the FRN (Hauser et al., 2014). Using single-trial analysis of the EEG, researchers found that prediction error resulting from perceptual salience (i.e., the phase coherence of face and house images) elicited feedback-evoked FRN (Lou et al., 2015). The P300 is often elicited by rare or novel stimuli. The P300 may reflect unexpected changes in the sensory environment that are sufficiently salient to enter the awareness (Pinheiro, Barros, & Pedrosa, 2015). Multi-modal neuroimaging research showed that the P300 was significantly correlated with the activity in the striatum (Pfabigan et al., 2014; Pogarell et al., 2011). Taken together, the abovementioned neuroimaging and electrophysiology studies have consistently indicated a role of the striatum in salience-based reward processing. Our present findings have corroborated the function of the striatum in integrating subjective value and visual salience, and further demonstrated the functional dissociations of the striatum subregions in salience-based specific outcome evaluation.

### The striatum and prefrontal cortex encode salience-modulated behavioral adjustment

Our fMRI results demonstrated a crucial role of the vmPFC in salience-guided strategic behavioral adjustment. Our study is thus among the first to identify a crucial role of vmPFC in salience-driven decision-making. These findings are consistent with the general evidence of vmPFC in the interplay of goal-directed and habitual learning (Piray et al., 2016), or a transition from goal-directed to habitual action control (Gremel & Costa, 2013). Neuroimaging studies and metaanalysis have suggested an important contribution of the prefrontal cortex in guiding goal-directed instrumental choice (Huang et al., 2020; Valentin, Dickinson, & O’Doherty, 2007), avoidance learning (Kim, Shimojo, & O’Doherty, 2006), and encoding abstract rules in complex decisions (J. P. O’doherty, Hampton, & Kim, 2007), which are consistent with our present findings of behavioral adjustment. Although the striatum and vmPFC have long been implicated in the value computation (Bartra et al., 2013; Rangel, Camerer, & Montague, 2008) and can be further modulated by the self-control or selective attention (Hare, Camerer, & Rangel, 2009; Hare, Malmaud, & Rangel, 2011), we here for the first time revealed a functional role of the vmPFC in salience-guided behavioral adjustment.

Moreover, high-order PPI analysis suggested that activity in the vmPFC was functionally correlated with the NAcc (for utility emphasis) and dmPFC (for performance emphasis), and the strength of their connectivity can further explain the interindividual difference in switching frequency after salience manipulation, respectively. However, the vmPFC and dmPFC showed a difference in encoding feedback-guided behavioral adjustments, in which the vmPFC encoded both utility- and performance-guided behavioral adjustments and was further modulated by salience manipulation, while dmPFC only encoded performance-guided behavioral adjustment after salience manipulation. These findings suggested that the role of the vmPFC in salience-modulated decision-making could be feedback general (both utility and performance), while the dmPFC could be feedback-specific (performance only), comparable with the findings of the dorsal striatum in feedback-general outcome evaluation while the ventral striatum only in performance-based outcome evaluation. Altogether, our results demonstrated a crucial role of the frontal-striatal circuit in encoding salience-driven valuation and behavioral adjustments, pointing that salience-driven process may integrate both top-down goal-directed and bottom-up habitual learning processes (Gremel & Costa, 2013; Huang et al., 2020; Piray et al., 2016; Redgrave et al., 2010).

Lastly, the ventral striatum was involved in both salience-modulated outcome evaluation (mainly the putamen) and behavioral adjustment (mainly the NAcc), whereas the dorsal striatum (i.e., caudate) was only engaged in salience-modulated outcome evaluation, further demonstrating a functional dissociation of the striatum subregions in discriminating decision stages (i.e., outcome evaluation or action), that were consistent with the crucial role of the ventral striatum in value integration (Bartra et al., 2013).

### Limitations and future directions

First, in our present study, there was no inherent statistical structure to the task, and therefore participants could not learn the distribution of outcomes from feedback. Such a design may exacerbate the salience-driven incidental learning, in which the participants only adjusted their decisions along with the salient dimension and the feedback from the most immediate trial. Although our findings may explain how salience exerts influence on the attention and subjective valuation systems that further strengthen incidental learning, it is still unclear where the default incidental learning comes from. Further studies may answer these questions independently under a non-salience condition. Second, although we have measured the subjective satisfaction ratings towards outcomes, it is still unclear whether and how attention, motivation, and emotion (i.e., rejoice and disappointment) may interplay with stable personality traits that further determine the individual difference. Future studies may break down the potential factors in giving rise to behavioral differences in learning. Last, in our present study, visual salience exerted its effect by explicitly instructing participants to pay attention to certain aspects of reward feedback. The salience cues were thus not exogenous because specific verbal or non-verbal instructions had been provided to them; and it was, therefore, motivational salience by virtue of the intrinsic properties or behavioral importance as mentioned by Zink et al (Zink et al., 2004). Salience can also be manipulated by other means, such as the amount of time participants fixate on an item (Armel et al., 2008), the amount of information revealed to participants (Frydman & Rangel, 2014), or visual contrasts (Moher, Anderson, & Song, 2015). It remains to be tested whether our findings can be extended to settings using endogenous salience, and how our findings may guide irrational (i.e., overspending) or addictive (i.e., food, tobacco addiction) behavior intervention via explicit (i.e., emphasis) and implicit (i.e., neurofeedback) attention manipulation techniques.

### Conclusion

In conclusion, our behavioral studies have revealed that specific visual salience modulates behavioral strategies based on feedback evaluation. Our eye-tracking studies have further established a crucial role of attention in salience-driven outcome evaluation, that further guides incidental learning. Lastly, our fMRI results identified the neural correlates of salience-modulated incidental learning, especially the role of the striatum and vmPFC in encoding salience-modulated outcome evaluation and behavioral adjustment. Our findings suggest that the reward system is orchestrated by visual salience and highlight the key role of attention and the frontal-striatal circuit in visual salience-guided incidental learning and decision biases. Such salience modulation can be utilized in real-life situations, such as casino, marketing, and policy making, to nudge individuals’ choices.

## Acknowledgments

We thank Ping Zhang and Shuyi Wu for assisting with the fMRI data collection, and Chuhua Cai for collecting all the eye-tracking data. This research was supported by the Tohoku University Operational Budget of President’s Discretionary Funds (Research) (No. 56045090) and Japan Society for the Promotion of Science Grant-in-Aid for Early-Career Scientists (No.22K15626) (to S.S.) and the NSF (BCS-1945230) and the Dana Foundation (to S.W.). The funders had no role in the study design, data collection, analysis, decision to publish, or preparation of the manuscript.

## CRediT Statement

Sai Sun: Conception, Data curation, Formal analysis, Methodology, Visualization, Writing— original draft, Writing—review and editing. Hongbo Yu: Writing—original draft, Writing— review and editing. Shuo Wang: Conception, Methodology, Visualization, Writing—original draft, Writing—review and editing, Supervision. Rongjun Yu: Conception, Methodology, Visualization, Writing—original draft, Writing—review and editing, Supervision. All authors approved the final version of the manuscript for submission. The authors have no conflicts of interest to declare.

## Data and code availability statement

The data and code used in this study is publicly available on OSF (https://osf.io/ud7yc/).

## Supplemental Figure Legends

**Fig. S1.** Absolute switching frequency for each condition. Legend conventions as Fig. 2 and Fig. 4. **(A, B)** Exp.2, specific salience: behavioral study. **(C, D)** Exp.3, non-specific salience: behavioral study. **(E, F)** Exp.4, specific salience: eye-tracking study. **(G, H)** Exp.5: non-specific salience: eye-tracking study. **(I, J)** Exp.6, specific salience: fMRI study. **(A, C, E, G, I)** Emphasis on utility. **(B, D, F, H, J)** Emphasis on performance. **(A, B, E, F, I, J)**. Specific emphasis. **(C, D, G, H)**. Non-specific emphasis. WC: win-correct. WI: win-incorrect. LC: loss-correct. LI: loss-incorrect. Error bars denote one SEM across participants. Asterisk indicates a significant difference using two-tailed one-sample t-test: +: P < 0.1, *: P < 0.05, **: P < 0.01, and ***: P < 0.001. n.s.: nonsignificant. Red: congruent / salient. Gray: incongruent / non-salient. Solid bars denote loss-win whereas open bars denote incorrect-correct.

**Fig. S2.** Absolute fixation difference for each condition. Legend conventions as Fig. 3. (A1-B4) Specific salience. **(A1-A3)** Emphasis on utility. **(A1)** Total fixation for each condition; **(A2)** Fixation difference (Chosen-Unchosen) for each condition; **(A3)** Normalized fixation difference for each condition. **(B1-B3)** Emphasis on performance. **(B1)** Total fixation for each condition; **(B2)** Fixation difference (Chosen-Unchosen) for each condition; **(B3)** Normalized fixation difference for each condition. **(A4)** Salience-guided fixation difference; **(B4)** Salience-guided normalized fixation difference. **(C1-D4)** Non-specific emphasis. **(C1-C3)** General emphasis on utility. **(C1)** Total fixation for each condition; **(C2)** Fixation difference (Chosen-Unchosen) for each condition; **(C3)** Normalized fixation difference for each condition. **(D1-D3)** General emphasis on performance. **(D1)** Total fixation for each condition; **(D2)** Fixation difference (Chosen-Unchosen) for each condition; **(D3)** Normalized fixation difference for each condition. **(C4)** Non-specific salience-guided fixation difference; **(D4)** Non-specific salience-guided normalized fixation difference. WC: win-correct. WI: win-incorrect. LC: loss-correct. LI: loss-incorrect. Error bars denote one SEM across participants. Asterisk indicates a significant difference using two-tailed one-sample t-test: +: P < 0.1, *: P < 0.05, **: P < 0.01, and ***: P < 0.001. n.s.: non-significant. Red: congruent / salient. Gray: incongruent / non-salient. Solid bars denote loss-win whereas open bars denote incorrect-correct.

**Fig. S3.** Response time (RT) of staying or switching choices for each condition. **(A1-B3)** Specific salience. RT for behavioral switching when emphasis on utility **(A1)** and performance **(A2)**. RT for behavioral staying when emphasis on utility **(B1)** and performance **(B2)**. Salience-guided behavioral switching **(A3)** and behavioral staying **(B3)**. **(C1-D3)** Non-specific salience. RT for behavioral switching during general emphasis on utility **(C1)** and performance **(C2)**. RT for behavioral staying during general emphasis on utility **(D1)** and performance **(D2)**. Non-specific salience-guided behavioral switching **(C3)** and behavioral staying **(D3)**. WC: win-correct. WI: win-incorrect. LC: loss-correct. LI: loss-incorrect. Error bars denote one SEM across participants. Asterisk indicates a significant difference using two-tailed one-sample t-test: +: P < 0.1, *: P < 0.05, **: P < 0.01, and ***: P < 0.001. n.s.: non-significant. Red: congruent / salient. Gray: incongruent / non-salient. Solid bars denote loss-win whereas open bars denote incorrect-correct.

**Fig. S4.** Correlation between attention and behavioral switching. (**A1-A4**) Specific emphasis. **(A1, A2)** Specific emphasis on utility. **(A1)** Utility-based behavioral switching was positively correlated with utility-based fixation difference; **(A2)** Performance-based behavioral switching was positively correlated with performance-based fixation difference. **(A3, A4)** Specific emphasis on performance. **(A3)** Utility-based behavioral switching was not correlated with utility-based fixation difference. **(A4)** Performance-based behavioral switching was positively correlated with performance-based fixation difference. (**B1-B4**) Non-specific emphasis. **(B1, B2)** Non-specific emphasis on utility dimension. No significant correlation was observed between attention and behavioral switching. **(B3, B4)** Non-specific emphasis on performance dimension. No significant correlation was observed between attention and behavioral switching. **(C1, C2)** Specific emphasis. Chosen value, unchosen value, and normalized fixation difference, significantly contributed to the behavioral strategy when either utility **(C1)** or performance **(C2)** was emphasized. **(D1, D2)** Nonspecific emphasis. Chosen value and unchosen value significantly contributed to the behavioral strategy when either utility **(D1)** or performance **(D2)** was emphasized. However, normalized fixation difference didn’t play a significant role in predicting behavioral strategy when either utility (**D1**) or performance **(D2)** was emphasized. *: P < 0.05, **: P < 0.01, and n.s.: non-significant.

**Fig. S5.** Control results for salience-modulated outcome evaluation in fMRI study. Legend conventions as **Fig. 5**. It is worth noting that the value of the chosen card (i.e., ‘chosen value’) parametrically reflected win-loss information, whereas the difference in values between the chosen and unchosen cards (i.e., ‘chosen-unchosen’ / relative value) parametrically reflected correct-incorrect information.

