## Supplementary figures and images for "Cognitive and neural bases of salience-driven incidental learning"

### Supplemental Figure 1

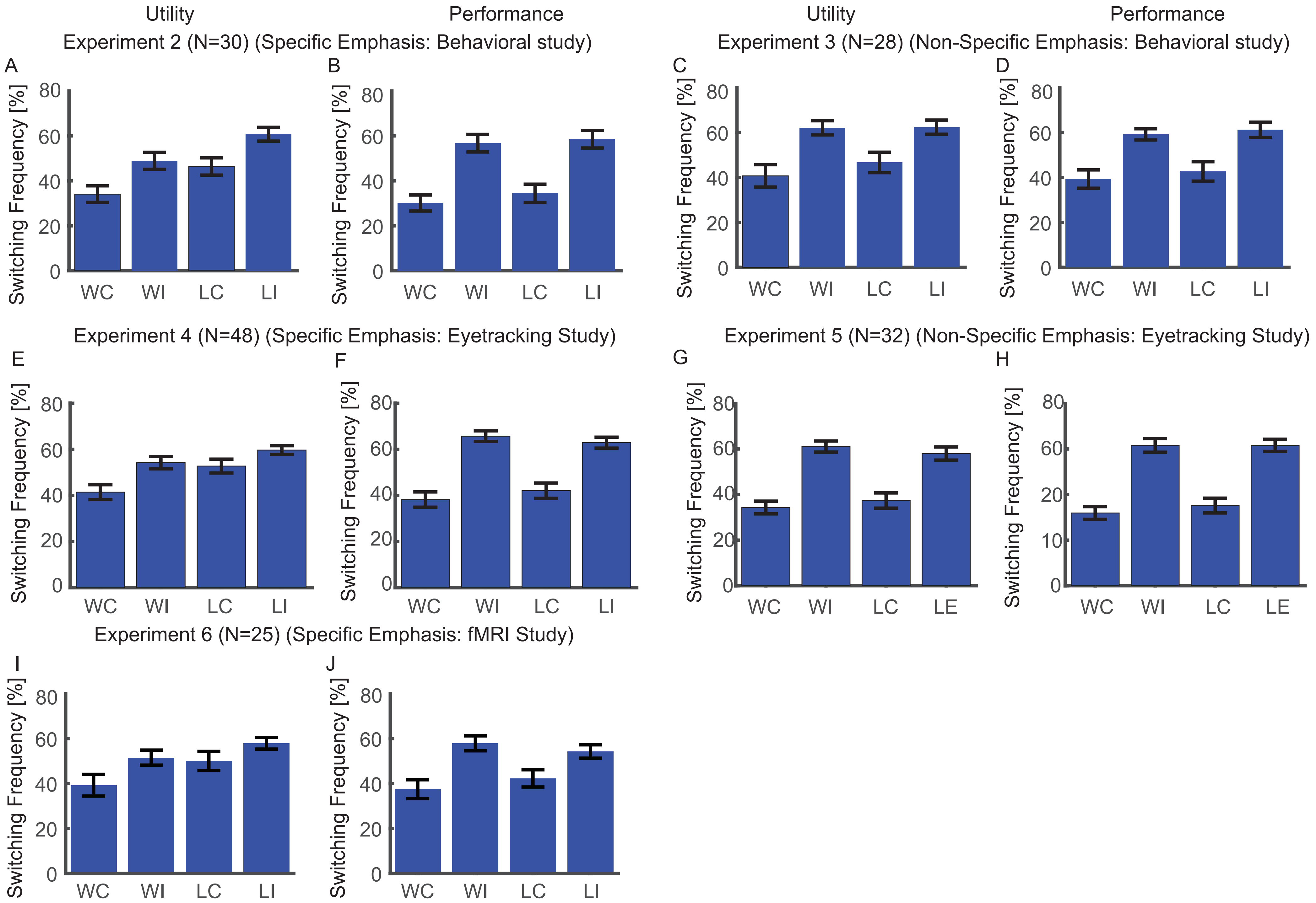

### Supplemental Figure 2

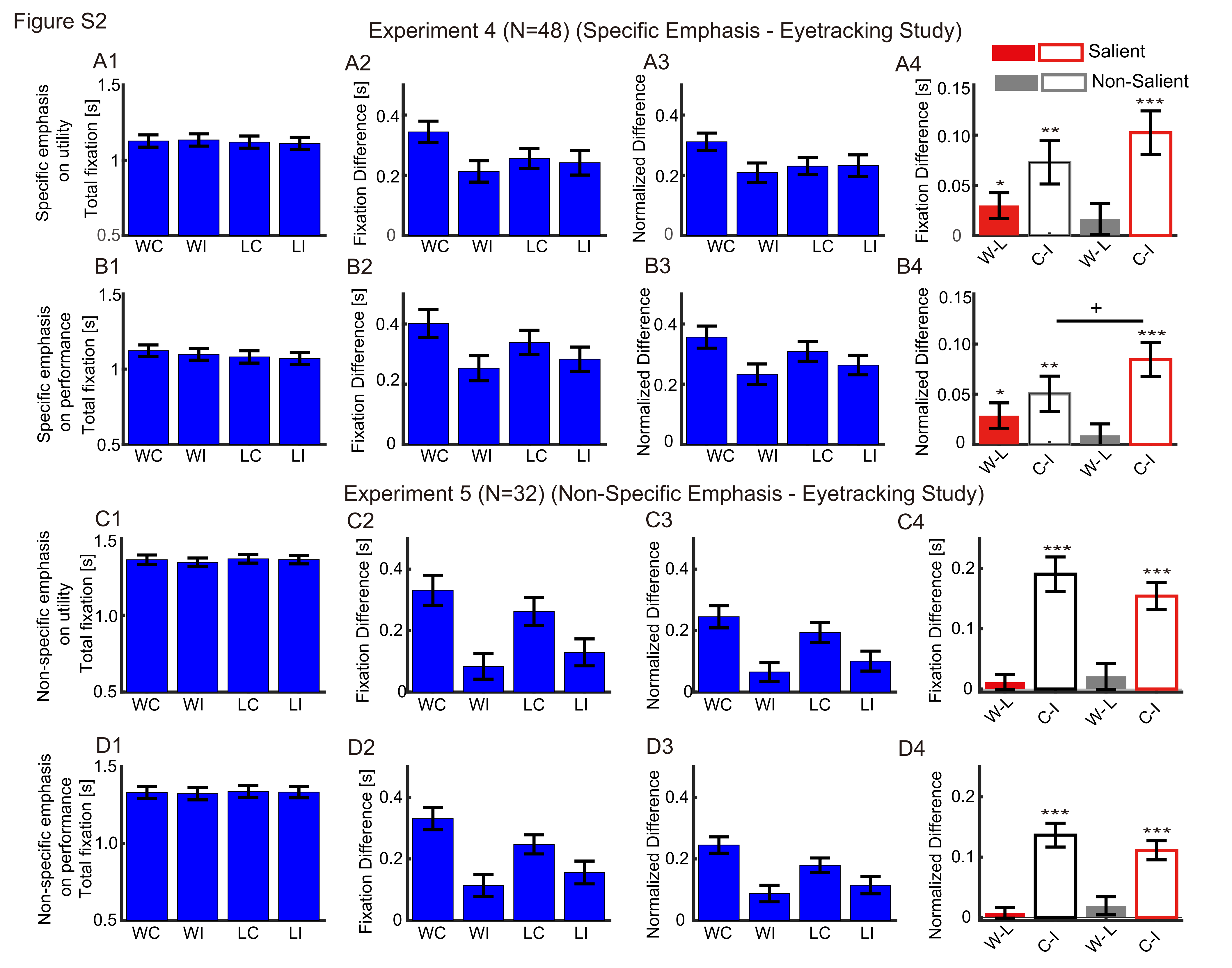

### Supplemental Figure 3

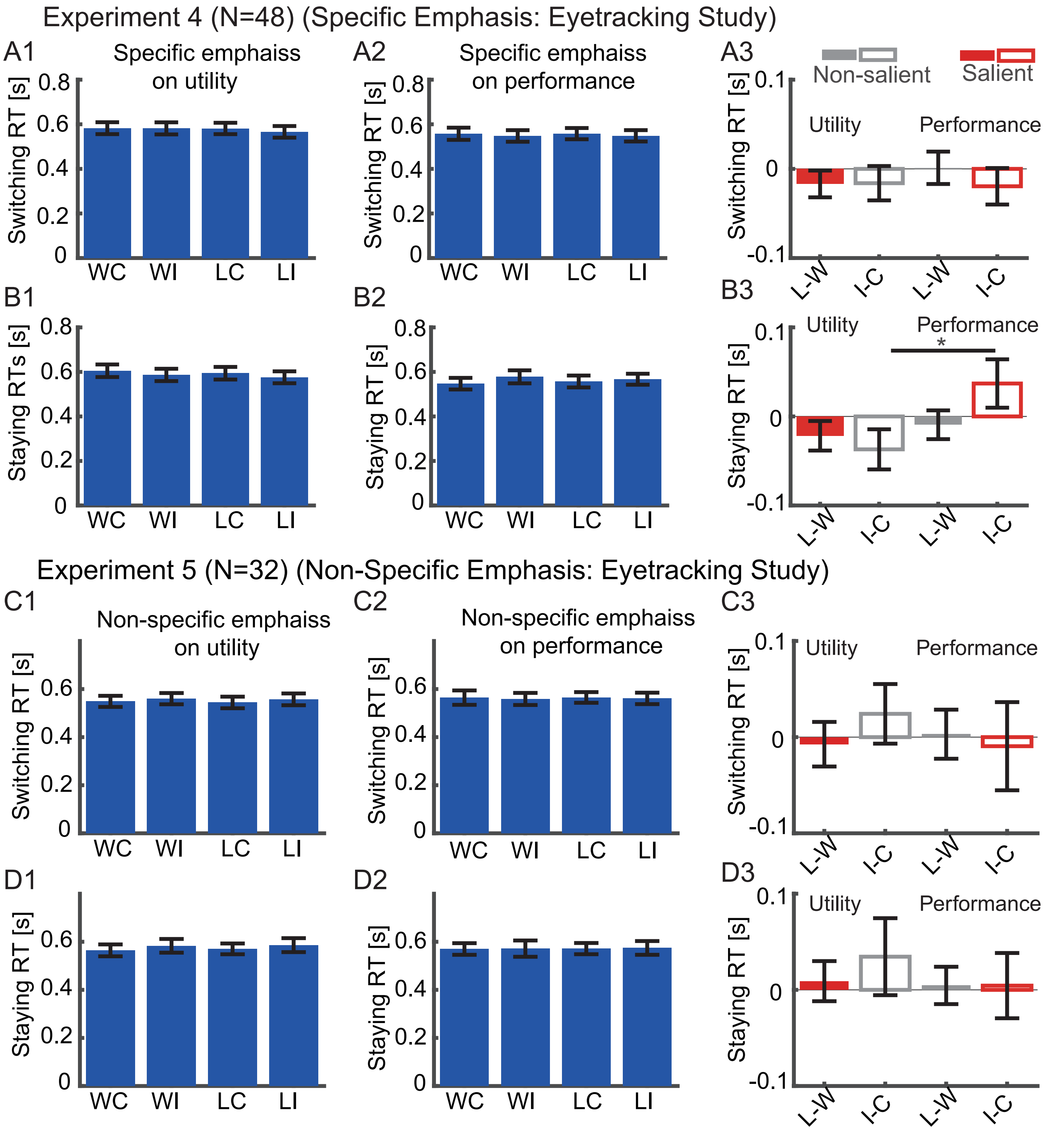

### Supplemental Figure 4

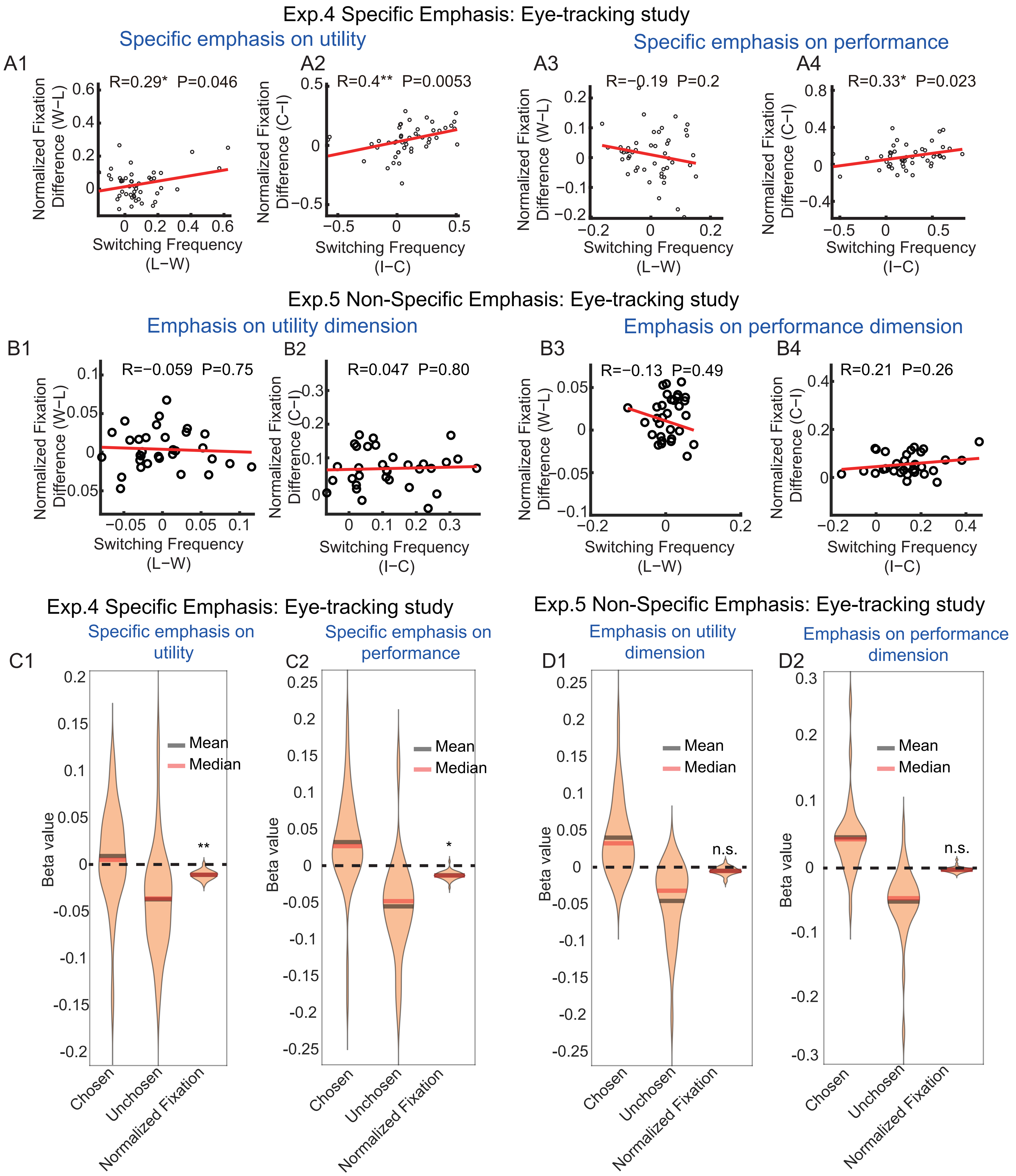

### Supplemental Figure 5

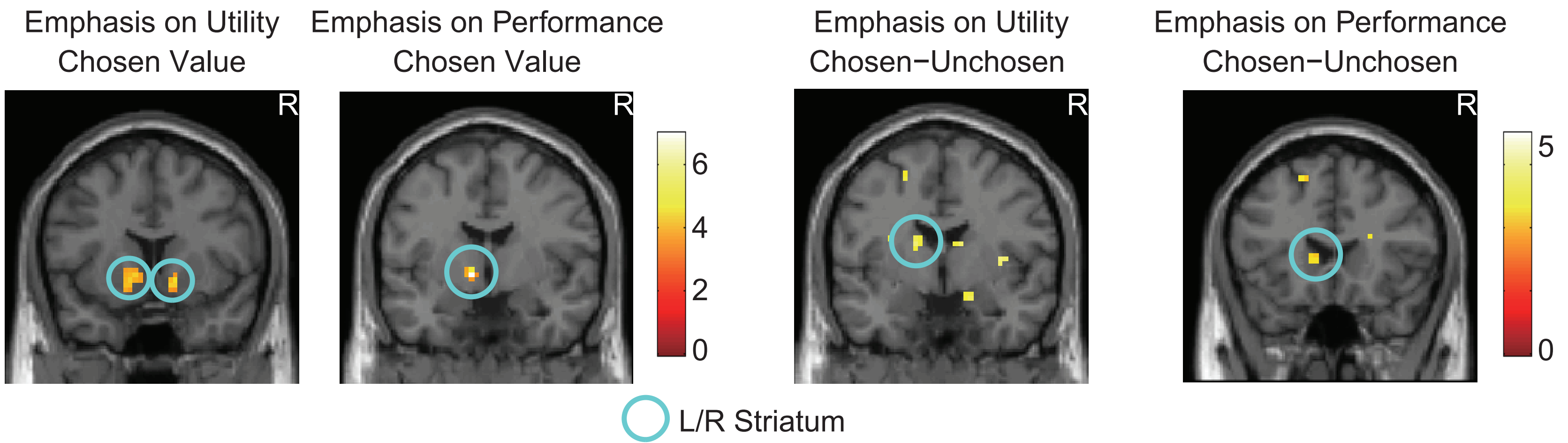
