## Supplemental Table 1 for "Cognitive and neural bases of salience-driven incidental learning"

**Table S1.** Brain areas are modulated by salience-modulated outcome evaluation and behavioral adjustment. All values are P < 0.001 uncorrected at the peak voxel level. * indicates P < 0.05 and ** indicates P < 0.01 family-wise error (FWE) after small volume correction (SVC).

<table>
<tr><th></th><th>Contrast</th><th>Brain Region</th><th>Z-score</th><th colspan="3">Peak Coordinate MNI (X Y Z)</th><th>Volume (voxel)</th></tr>
<tr><td colspan="8">Salience-modulated outcome evaluation</td></tr>
<tr><td rowspan="10">Emphasis on utility</td><td rowspan="2">Win−Loss</td><td>Striatal region<br>L Caudate head/nucleus**</td><td>4.04**</td><td>−9</td><td>12</td><td>0</td><td>23</td></tr>
<tr><td>L Occipital Lobe</td><td>4.44</td><td>−30</td><td>−87</td><td>−9</td><td>130</td></tr>
<tr><td rowspan="8">Correct−Incorrect</td><td>Striatal regions<br>L Caudate/Caudate head/Caudate nucleus**<br>L Caudate body**<br>L Putamen**<br>L Nucleus Accumbens**<br>R Caudate**<br>R Putamen**<br>R Medial Global Pullidus*<br>R Nucleus Accumbens**</td><td>5.15**<br>4.55**<br>5.19**<br>5.23**<br>5.21**<br>4.47**<br>3.70*<br>4.52**</td><td>−12<br>−12<br>−15<br>−18<br>9<br>15<br>12<br>12</td><td>18<br>18<br>9<br>6<br>18<br>9<br>3<br>6</td><td>6<br>9<br>−9<br>−15<br>9<br>−9<br>−6<br>−15</td><td>107<br>22<br>186<br>16<br>178<br>25<br>2<br>20</td></tr>
<tr><td>L Precuneus</td><td>4.68</td><td>−15</td><td>−57</td><td>36</td><td>457</td></tr>
<tr><td>L/R Posterior Cingulate Cortex</td><td>4.04<br>3.59</td><td>−12<br>6</td><td>−51<br>−45</td><td>21<br>21</td><td>104</td></tr>
<tr><td>Middle Cingulate Cortex</td><td>4.17</td><td>0</td><td>−39</td><td>39</td><td>71</td></tr>
<tr><td>L/R Anterior Cingulate Cortex</td><td>3.67<br>3.93</td><td>−9<br>15</td><td>45<br>39</td><td>−3<br>6</td><td>25<br>63</td></tr>
<tr><td>L vmPFC</td><td>3.65</td><td>−6</td><td>57</td><td>9</td><td>55</td></tr>
<tr><td>L/R Occipital cortex</td><td>4.59</td><td>36<br>−21<br>−6</td><td>−69<br>−99<br>−99</td><td>3<br>18<br>18</td><td>577</td></tr>
<tr><td>L OFC/BA11</td><td>4.47</td><td>−36</td><td>39</td><td>−9</td><td>169</td></tr>
<tr><td>Emphasis on performance</td><td>Win−Loss</td><td>Striatal region<br>L Lateral/Medial Globus Pallidus**</td><td>4.20**</td><td>−9</td><td>−3</td><td>−3</td><td>13</td></tr>
</table>

| | | | | | | | |
|---|---|---|---|---|---|---|---|
| | Correct−Incorrect | Striatal region<br>L/R Caudate<br>L/R Putamen<br>L/R Nucleus Accumbens | <br>6.67<br>6.65<br>6.30 | <br>−15<br>−18<br>−9 | <br>6<br>12<br>21 | <br>−15<br>−6<br>3 | 676 |
| | | L PCC/Precuneus | 6.27 | −3<br>0 | −45<br>−24 | 39<br>51 | 894 |
| | | L Hippocampus** | 5.98** | −33 | −6 | −27 | 21 |
| | | L/Middle vACC | 5.92 | −6<br>−3<br>0 | 51<br>48<br>42 | 6<br>−9<br>3 | 206 |
| **Salience-modulated behavioral adjustment** | | | | | | | |
| Emphasis on utility | Switch(L−W)<br>−Stay(L−W) | L vACC/vmPFC<br>L vACC* | 3.74<br>3.73* | −12<br>−12 | 39<br>42 | 0<br>0 | 32<br>12 |
| | Switch(I−C)<br>−Stay(I−C) | No significant activation | | | | | |
| Emphasis on performance | Switch(L−W)<br>−Stay(L−W) | R Nucleus Accumbens *<br>L Temporal lobe | 2.96*<br>4.08 | 12<br>−51 | 6<br>−39 | −12<br>15 | 2<br>76 |
| | Switch(I−C)<br>−Stay(I−C) | vmPFC<br>L vmPFC**<br>R vmPFC**<br>L vACC* | 3.73<br>3.50**<br>3.70**<br>3.18* | 0<br>−3<br>3<br>−9 | 54<br>54<br>54<br>42 | 3<br>3<br>3<br>6 | 269<br>64<br>64<br>41 |
