## Supplemental Table 2 for "Cognitive and neural bases of salience-driven incidental learning"

**Table S2.** Brain areas are modulated by chosen or relative values. All values are P < 0.005 uncorrected. It is worth noting that the value of the chosen card (i.e., ‘chosen value’) parametrically reflected win−loss information, whereas the difference in values between the chosen and unchosen cards (i.e., ‘chosen−unchosen ’/ relative value) parametrically reflected correct−incorrect information.

| Conditions | Contrast | Brain Region | Z-score | Peak Coordinate MNI (X Y Z) | | | Volume (voxel) |
|---|---|---|---|---|---|---|---|
| Emphasis on utility | Absolute value (Chosen value) | L/R caudate* | 3.15<br>3.14* | −9<br>12 | 9<br>9 | −3<br>−9 | 38<br>13 |
| | | L medial global pullidus* | 2.58* | −15 | −3 | −3 | 1 |
| | | L/R putamen* | 3.17*<br>3.26* | −30<br>15 | 0<br>9 | 6<br>−6 | 3<br>12 |
| | | L/R nucleus accumbens** | 3.11**<br>3.28** | −12<br>12 | 6<br>6 | −9<br>−12 | 5<br>10 |
| | Relative value (Chosen−Unchosen) | L caudate head* | 2.81* | −12 | 21 | 3 | 2 |
| Emphasis on performance | Absolute value (Chosen value) | L medial global pullidus* | 3.20*<br>4.77 | −12<br>−9 | 0<br>−3 | −3<br>−3 | 2<br>17 |
| | Relative value (Chosen-Unchosen) | L caudate head* | 3.74* | −12 | 21 | 3 | 2 |
